# Structural basis of *Mycobacterium tuberculosis* transcription and transcription inhibition

**DOI:** 10.1101/099606

**Authors:** Wei Lin, Soma Mandal, David Degen, Yu Liu, Yon W. Ebright, Shengjian Li, Yu Feng, Yu Zhang, Sukhendu Mandal, Yi Jiang, Shuang Liu, Matthew Gigliotti, Meliza Talaue, Nancy Connell, Kalyan Das, Eddy Arnold, Richard H. Ebright

## Abstract

**One Sentence Summary:** Structures of *Mycobacterium tuberculosis* RNA polymerase reveal taxon-specific properties and binding sites of known and new antituberculosis agents

**Abstract:** *Mycobacterium tuberculosis* (*Mtb*) is the causative agent of tuberculosis, which kills 1.8 million annually. *Mtb* RNA polymerase (RNAP) is the target of the first-line antituberculosis drug rifampin (Rif). We report crystal structures of *Mtb* RNAP, alone and in complex with Rif. The results identify an *Mtb*-specific structural module of *Mtb* RNAP and establish that Rif functions by a steric-occlusion mechanism that prevents extension of RNA. We also report novel non-Rif-related compounds–Nα-aroyl-N-aryl-phenylalaninamides (AAPs)–that potently and selectively inhibit *Mtb* RNAP and *Mtb* growth, and we report crystal structures of *Mtb* RNAP in complex with AAPs. AAPs bind to a different site on *Mtb* RNAP than Rif, exhibit no cross-resistance with Rif, function additively when co-administered with Rif, and suppress resistance emergence when co-administered with Rif.

---

Rifampin (Rif) is the cornerstone of current antituberculosis therapy (*1-3*). The emergence and spread of Rif-resistant *Mycobacterium tuberculosis* (*Mtb*) is an urgent public-health crisis (0.6 million new cases annually; *1-3*). Rif-resistance in *Mtb* arises from substitution of residues of the binding site for Rif on its molecular target, *Mtb* RNA polymerase (RNAP; *2-3*). Intensive efforts are underway to identify Rif derivatives that are unaffected by substitutions in the Rif binding site, and to identify novel, non-Rif-related RNAP inhibitors that function through binding sites on RNAP that do not overlap the Rif binding site and thus are unaffected by substitutions in the Rif binding site (*3-4*). However, these efforts have been hampered by the unavailability of structural information for *Mtb* RNAP or a closely related bacterial RNAP. [Crystal structures of bacterial RNAP hitherto have been available only for *Thermus aquaticus* (*Taq*), *Thermus thermophilus* (*Tth*), and *Escherichia coli* (*Eco*) RNAP, which have ≤40% sequence identity with *Mtb* RNAP (*5*), although a paper reporting a structure of *Mycobacterium smegmatis* RNAP is in press (*6*).]

Here, we have determined crystal structures of *Mtb* RNAP transcription initiation complexes at 3.6-4.4 Å resolution using a strategy analogous to the strategy used to determine structures of *Tth* RNAP transcription initiation complexes (*7*): i.e., crystallization of *Mtb* RNAP σ^A^ holoenzyme in complex with a synthetic nucleic-acid scaffold that mimics the ssDNA transcription bubble and downstream dsDNA of a catalytically competent RNAP promoter open complex (RPo), or with the same synthetic nucleic-acid scaffold in the presence of synthetic RNA oligomers corresponding to 2, 3, and 4 nt RNA products (RNAP-promoter initial transcribing complexes, RPitc2, RPitc3, and RPitc4; Tables S1-S3; Figs. S1-S2).

The resulting structures of *Mtb* RPo and RPitc are similar to previously reported structures of *Tth* RPo and RPitc (*7*) in overall structural organization; in sequence-specific interactions between RNAP holoenzyme and promoter −10, discriminator, and core recognition elements; and in sequence-independent interactions between RNAP holoenzyme and template-strand ssDNA and downstream dsDNA (Figs. 1A, S3-S6).

**Fig. 1.**
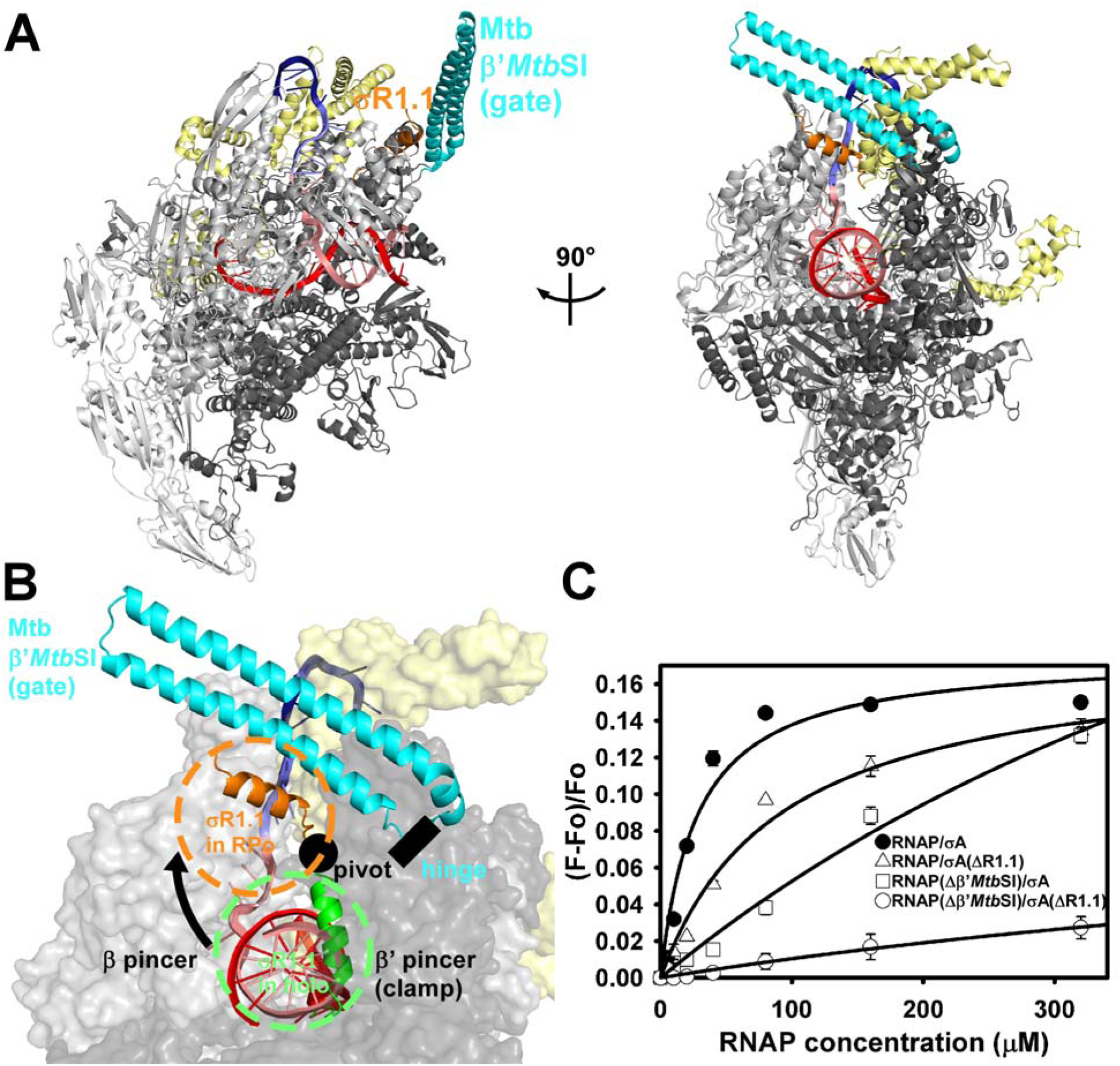
Structure of *Mtb* RPo. (**A**) Overall structure of *Mtb* RPo (two orthogonal views). Gray, RNAP; yellow, σ^A^; blue, −10 element of DNA nontemplate strand; light blue, discriminator element of DNA nontemplate strand; pink, rest of DNA nontemplate strand; red, DNA template strand; cyan, taxon-specific sequence insertion (β’*Mtb*SI); orange, σR1.1 H4. (**B**) Trapping of dsDNA in RNAP active-center cleft by β’*>Mtb*SI and σR1.1. Green and orange a-helices, σR1.1 in *Eco* RNA holoenzyme and *Mtb* RPo; green and orange dashed circles, approximate molecular volumes of σR1.1 in *Eco* RNAP holoenzyme and *Mtb* RPo. Other colors as in (A). (**C**) Effects of deletion of β’*Mtb*SI, σR1.1, or both on stability of *Mtb* RPo.

The structures reveal two novel features.

First, the structures reveal the conformation and interactions of a ~100 residue taxon-specific sequence insertion present in the β′ subunit of RNAP from Mycobacterial and closely related Acinetobacterial taxa (β’*Mtb*SI; β’In1 in *8*; β′ residues Fig. 1A). The structures of *Mtb* RPo and RPitc show that β’*Mtb*SI folds as an extraordinarily long (~70 Å) α-helical coiled-coil (Figs. 1A, S2), and a structure at 2.2 Å resolution of an isolated protein fragment corresponding to β’*Mtb*SI shows a superimposable conformation and thus shows that β’*Mtb*SI folds independently (Fig. S7; Table S3). In the structures of *Mtb* RPo and RPitc, the β’*Mtb*SI coiled-coil emerges from the RNAP clamp module and extends across the RNAP active-center cleft. We infer that the β’*Mtb*SI coiled-coil serves as a “gate” that helps trap and secure DNA within the active-center cleft. and that presumably must open, rotating about a “hinge” formed by the short unstructured segments connecting β’*Mtb*SI to the remainder of β’, to allow DNA to enter the active-center cleft (Figs. 1A-B, S2). Consistent with the inference from the structures that β’*Mtb*SI helps trap and secure DNA within the active-center cleft, deletion of β’*Mtb*SI strongly reduces the stability of *Mtb* RPo (30-fold defect; Fig. 1C).

Second, the structures provide information about the position of σR1.1, a σ module that, based on fluorescence resonance energy transfer (FRET) data, occupies the dsDNA binding site within the RNAP active-center cleft in RNAP holoenzyme but is displaced from the dsDNA binding site upon formation of RPo (*9*; Fig. 1A-B). A structure of *Eco* RNAP holoenzyme shows that σR1.1 occupies the dsDNA binding site for dsDNA in RNAP holoenzyme (*10*), consistent with the FRET data (*9*), but previous structures of bacterial RPo and RPitc have not resolved σR1.1 (*7*). The structures of *Mtb* RPo and RPitc show clear, unambiguous electron density for one α-helix of σR1.1, the α-helix that connects σR1.1 to the remainder of σ (H4, numbered as in 7; Fig. 1A-B). Strikingly, whereas in *Eco* RNAP, holoenzyme H4 is oriented perpendicular to the floor of the dsDNA binding site and occupies the dsDNA binding site (green in Figs. 1B, S8A), in *Mtb* RPo and RPitc, H4 is rotated by ~100°, about a “pivot” formed by the short unstructured segment between σR1.1 and the remainder of σ, and is oriented essentially parallel to the floor of the dsDNA binding site, outside the dsDNA binding site in *Mtb* RPo and RPitc (orange in Figs. 1B, S8A). The rotation of H4 in RPo and RPitc displaces the center of H4 by ~30 Â from its position in RNAP holoenzyme, consistent with the FRET data (*9*), and positions H4 between β’MtbSI and dsDNA (Figs. 1B, S8A). Assuming *Mtb* σR1.1 has approximately the same molecular volume as *Eco* σR1.1, σR1.1 in *Mtb* RPo and RPitc occupies essentially the entire space between β’*Mtb*SI and dsDNA, potentially making a continuous chain of β’*Mtb*SI-σR1.1-dsDNA interactions that close the active center and help trap and secure dsDNA in the active center (Figs. 1B, S8B). Consistent with this inference, deletion of *Mtb* σR1.1 strongly reduces the stability of *Mtb* RPo (5-fold defect), and deletion of both *Mtb* σR1.1 and β’*Mtb*SI very strongly reduces the stability of RPo (40-fold defect; Fig. 1C).

We determined a crystal structure of *Mtb* RPo in complex with Rif using analogous procedures (Fig. 2A-C; Table S2). The structure shows RNAP-Rif interactions similar to those in previously reported structures of *Tth* RNAP and *Eco* RNAP in complex with rifamycins (11-13), but, in this case, with a Rif binding-site sequence from a bacterial species for which Rif is a clinically relevant antibacterial drug (Fig. 2A-C). The structure shows direct H-bonded contacts between Rif and two of the three residues most frequently substituted in Rif-resistant *Mtb* clinical isolates (β H526 and S531) and direct van der Waals interactions between Rif and the third (β D516) (*2-3*; Fig. 2B-C).

**Fig. 2.**
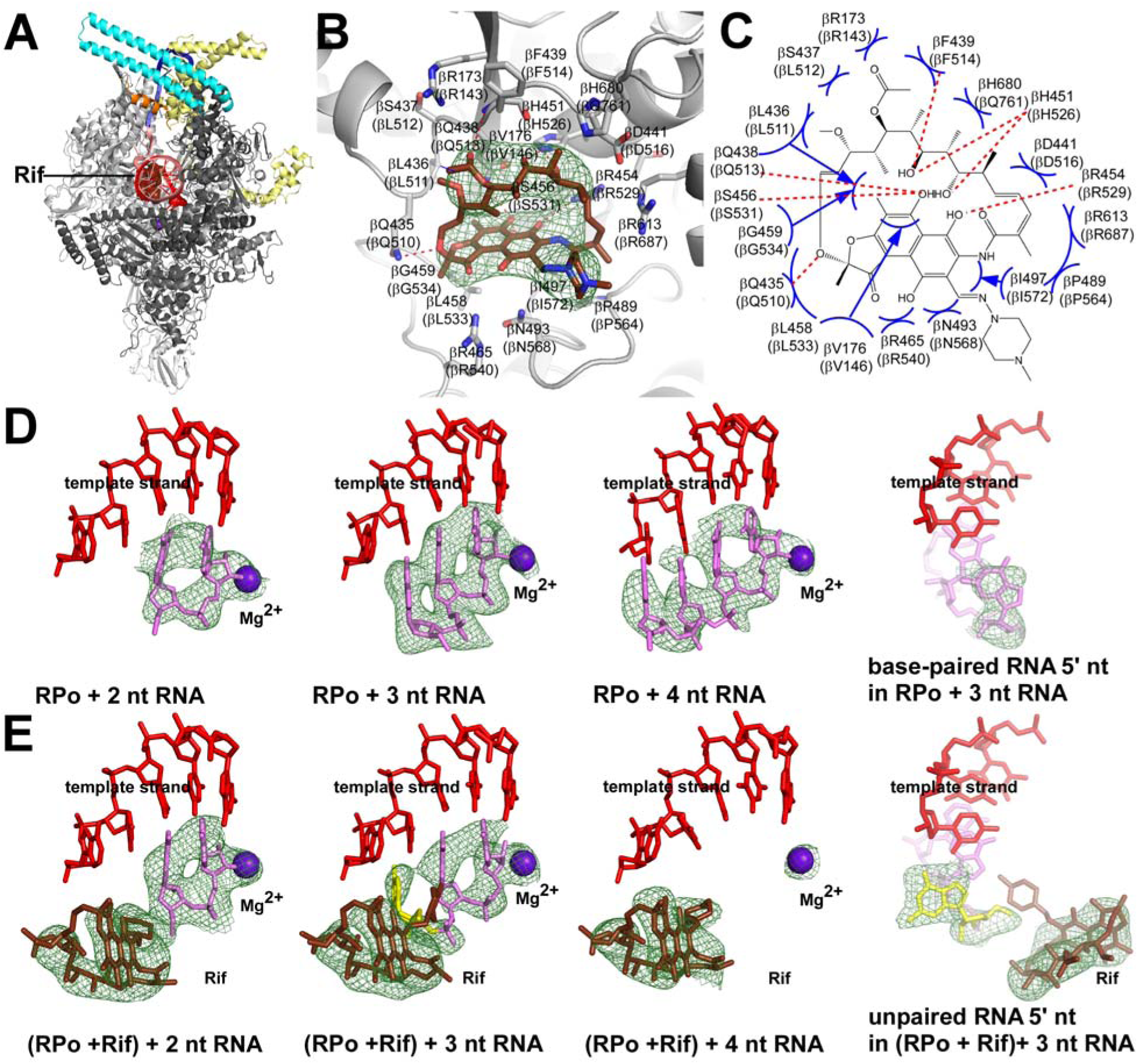
Structural basis of *Mtb* RNAP inhibition by Rif. (**A**) Overall structure of *Mtb* RPo in complex with Rif. Brown, Rif. Other colors as in Fig. 1. (**B**) *Mtb* RNAP-Rif interactions. Gray ribbons, RNAP backbone; gray and brown sticks, RNAP and Rif carbon atoms; red and blue sticks, RNAP and Rif oxygen and nitrogen atoms; green mesh, mFo-Fc electron density omit map for Rif (contoured at 2.5σ). Residues numbered as in *Mtb* RNAP and, in parentheses, as in *Eco* RNAP. (**C**) Summary of *Mtb* RNAP-Rif interactions. Red dashed lines, H-bonds; blue arcs, van der Waals interactions. (**D**)-(**E**) “Steric occlusion” mechanism for RNAP inhibition by Rif. Subpanels 1-3 of (D*),Mtb* RNAP crystallized with nucleic-acid scaffolds containing 2, 3, and 4 nt RNAs; subpanel 4 of (D), orthogonal view of subpanel 2 showing base pairing of RNA 5’ nucleotide. Subpanels 1-3 of (E*),Mtb* RNAP-Rif crystallized with nucleic-acid scaffolds containing 2, 3, and 4 nt RNAs; subpanel 4 of (E), orthogonal view of subpanel 2 showing unpaired, unstacked, RNA 5’ nucleotide. Red, DNA template strand; pink, RNA; yellow, unpaired, unstacked RNA 5’ nucleotide; brown, Rif; violet sphere, active-center catalytic Mg^2^+; green mesh, mFo-Fc electron density omit map for Rif, RNA, and catalytic Mg^2^+ (contoured at 2.5σ).

A series of crystal structures of *Mtb* RPitc in complex with Rif, obtained by crystallizing a pre-formed *Mtb* RNAP-Rif complex with nucleic-acid scaffolds containing RNA oligomers corresponding to 2, 3, and 4 nt RNA products (Table S3). The results graphically demonstrate that Rif inhibits transcription through a “steric occlusion” mechanism that prevents extension of 2-3 nt RNA products to yield longer RNA products–a mechanism that previously had been proposed (*11,14-15*), but that had not been directly demonstrated and had been controversial (*12,15*) (Fig. 2D-E; Tables S1-S2). Thus, whereas, in the absence of Rif, 2, 3, and 4 nt RNA products fully base pair to the DNA template strand, (Fig. 2D), in the presence of Rif, the 5′ nucleotide of a 3 nt RNA is unpaired, unstacked, and rotated by ~40°, due to steric clash with Rif (Fig. 2E, second and fourth panels), and a 4 nt RNA is unable to interact stably with the complex (i.e., shows no density; Fig. 2E, third panel). All the *Mtb* RPo-Rif and RPitc-Rif structures show clear, unambiguous electron density for the RNAP active-center catalytic Mg^2+^, inconsistent with the alternative, “allosteric” mechanism for Rif function proposed in *12* (Fig. 2D-E).

By high-throughput screening of a synthetic-compound library (details to be provided elsewhere), we have identified a novel class of small-molecule inhibitors of *Mtb* RNAP: Nɑ-aroyl-N-aryl-phenylalaninamides (AAPs; Fig. 3A). The prototype of the class, D-AAP1, exhibits potent, selective, stereospecific activity against Mycobacterial RNAP (potent inhibition of *Mtb* RNAP, but poor inhibition of other bacterial RNAP and human RNAP I, II, and III; Figs. 3A, S9) and exhibits potent, selective, stereospecific activity against Mycobacteria (potent activity against *Mtb, M. avium*, and *M. smegmatis*, but poor activity against other bacterial and mammalian cells; Figs. 3A, S9).

**Fig. 3.**
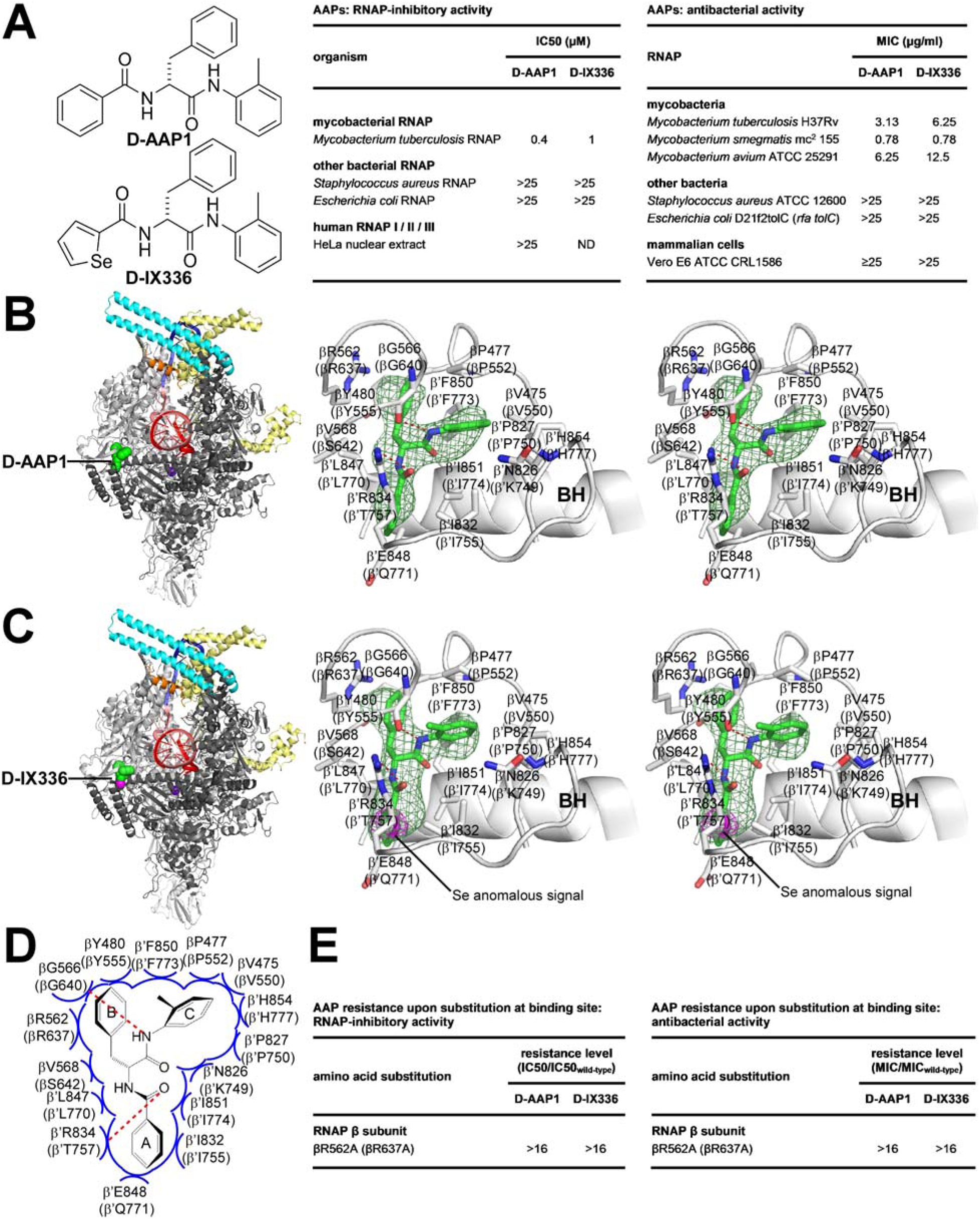
Structural basis of *Mtb* RNAP inhibition by AAPs. (**A**) Structures, RNAP-inhibitory activities, and antibacterial activities of D-AAP1 and selenium-containing analog D-IX336. (**B**)-(**C**) Structures of *Mtb* RPo in complex with D-AAP1 and D-IX336 (left panels, overall structures; right subpanels, stereoviews of *Mtb* RNAP-inhibitor interactions). Green surface, inhibitor; pink surface, selenium; green and pink sticks, inhibitor carbon and selenium atoms; pink mesh, selenium anomalous difference density (contoured at 4.5σ); BH, RNAP bridge helix. Other colors and labels as in Fig. 2B. (**D**) Summary of *Mtb* RNAP-inhibitor interactions. Colors and labels as in Fig. 2C. (**E**) Effects of Ala substitution of binding-site residue.

A crystal structure of *Mtb* RPo in complex with D-AAP1, determined by soaking preformed crystals of *Mtb* RPo with D-AAP1, defines the binding site, orientation, and interactions of D-AAP1 (Fig. 3B; Table S3). Synthesis of a D-AAP1 analog containing a carbon-to-selenium substitution, D-IX336, followed by crystal soaking, X-ray diffraction analysis, and selenium-anomalous-dispersion analysis confirms the identified binding site, orientations, and interactions (Figs. 3A-C, S10; Table S3). The structures reveal that AAPs bind to *Mtb* RNAP at a binding site centered on the N-terminus of the RNAP bridge helix (an ɑ-helix that bridges the RNAP active-center cleft and forms one wall of the RNAP active center; Fig. 3B-C). The three aromatic rings of the AAP bind in three pocket-like subsites (Figs. B-D). Alanine substitution of a residue of the observed binding site results in AAP-resistance, both for RNAP-inhibitory activity and for antibacterial activity, confirming the functional importance of the observed interactions (Fig. 3E). The structures enable rational, structure-based optimization of AAPs to improve potencies and properties. In particular, the structures show that the methyl group of ring “C” of an AAP projects into an unoccupied area (Figs. 3B-D), allowing substitution of this position with diverse chemical functionality. Lead-optimization efforts (to be described elsewhere) confirm the utility of the structures for structure-based lead optimization, and, in particular, confirm the ability to improve potency and properties by substitution of the methyl group of ring “C”.

The structures in Fig. 3 show that the binding site on RNAP for AAPs differs from, and does not overlap, the binding site on RNAP for Rif (Figs. 2A, 3B-C). The binding site on RNAP for AAPs is similar in location to the binding site for CBRs, a class of compounds that inhibit Gram-negative bacterial RNAP but do not inhibit Mycobacterial RNAP; *16-18*; Fig. S11). Thus, both AAPs (in *Mtb* RNAP) and CBRs (in Gram-negative RNAP) interact with the RNAP bridge-helix N-terminus (Figs. 3B-D; S11). We infer that AAPs are Mycobacteria-selective inhibitors that function through the bridge-helix N-terminus target, and CBRs are Gram-negative-selective inhibitors that function through the bridge-helix N-terminus. Comparison of structures of *Mtb* RNAP-AAP complexes and *Eco* RNAP-CBR complexes reveals the basis for the difference in selectivity: i.e., because of sequence differences, *Mtb* RNAP has a three-pocket site complementary to an AAP, with three rings, but *Eco* RNAP has a two-pocket site complementary to a CBR, with two rings (Figs. S11-S12). The crucial sequence differences apparently include β residue 642 and β′ residues 757 and 771, which line or approach the pocket that is present, and accommodates AAP ring “A”, in Mtb RNAP, but is absent in *Eco* RNAP (residues numbered as in *Eco* RNAP; Figs S11-S12). Based on the similarity in binding sites of AAPs and CBRs, AAPs most likely inhibit RNAP through a mechanism similar to that of CBRs: i.e., interference with bridge-helix conformational dynamics required for nucleotide addition (*17-18*).

A structure of *Mtb* RPo in complex with *both* D-AAP1 and Rif further confirms unequivocally that the AAP binding site differs from, and does not overlap, the Rif binding site and shows that an AAP and Rif can bind simultaneously to RNAP (Fig. 4A; Table S3).

**Fig. 4.**
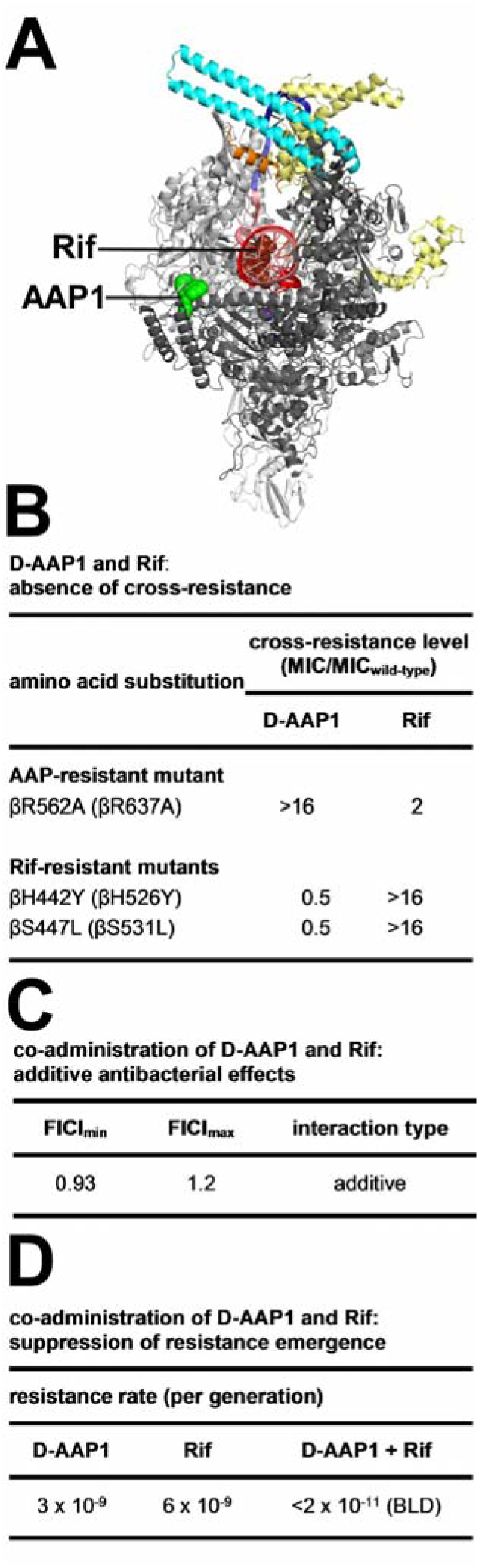
Additive antibacterial activity and suppressed resistance emergence upon co-administration of D-AAP1 and Rif. (**A**) Structure of *Mtb* RPo in complex with *both* D-AAP1 and Rif. Colors as in Figs. 2-3. (**B**) Absence of cross-resistance of D-AAP1 and Rif. (**C**) Additive antibacterial effects upon co-administration of D-AAP1 and Rif. (**D**) Suppressed resistance emergence upon co-administration of D-AAP1 and Rif.

The absence of overlap between the AAP and Rif binding sites (Fig. 4A) predicts that AAPs and Rif should not exhibit cross-resistance (since binding-site substitutions that interfere with binding of one compound should not affect binding of the other compound). Measurement of cross-resistance validates this prediction (Fig. 4B). Thus, a substitution in the AAP binding site confers AAP-resistance but not Rif-resistance, and substitutions in the Rif binding site confer Rif-resistance but not AAP-resistance (Fig. 4B).

The ability of AAPs and Rif to bind simultaneously to RNAP (Fig. 4A) predicts that co-administration of an AAP and Rif should result in additive antibacterial activity (since each compound should bind to its respective site and contribute to inhibition through its respective mechanism of inhibition). Measurement of inhibitor-inhibitor interactions in checkerboard interaction assays yields a fractional inhibitory concentration index (FICI) indicative of additive interactions, validating this prediction (Fig. 4C).

The absence of cross-resistance between AAPs and Rif (Fig 4B, together with the additive antibacterial activities of AAPs and Rif (Fig. 4C) predict that co-administration of an AAP and Rif should suppress resistance emergence (since resistance to co-administered AAP and Rif should require a rare double mutational hit inactivating two binding sites, rather than a single mutational hit inactivating one binding site). Luria-Delbrück fluctuation assays assessing spontaneous resistance rates for D-AAP1, Rif, and co-administered D-AAP1 and Rif, validate this prediction (Fig. 4D). Co-administering D-AAP1 and Rif results in a ≥ 100-fold reduction–to undetectable levels–in spontaneous resistance rates (Fig. 4D).

The structures presented here provide a foundation for understanding *Mtb* RNAP structure-function, for structure-based design of improved Rif derivatives effective against Rif-resistant *Mtb*, and for structure-based discovery and development of novel, non-Rif-related *Mtb* RNAP inhibitors effective against Rif-resistant *Mtb.* The novel, non-Rif-related *Mtb* RNAP inhibitors presented here–AAPs–exhibit potent antimycobacterial activity, no cross-resistance with Rif, additive antimycobacterial activity when co-administered with Rif, and suppression of resistance emergence–to undetectable levels–when co-administered with Rif. As such, AAPs represent *exceptionally promising* new lead compounds for development of new antituberculosis drugs.

## Acknowledgements

This work was supported by National Institutes of Health grants GM041376 and AI109713-8681 and Global Alliance for TB Drug Development contracts to R.H.E. We thank the Argonne National Laboratory for beamline access; S. Rodrigue, J. Mukhopadhyay, and C. Sassetti for plasmids; and S. Diamond, E. Lucumi, G. Waters, Z. Ma, K. Taneko, C. Cooper, K. Mdluli, and N. Fotouhi for discussion.

## Supplementary Materials

### Materials and Methods

#### Nα-benzoyl-N-(2-methylphenyl)-D/L-phenylalaninamide (DL-AAP1)

**DL-AAP1.**
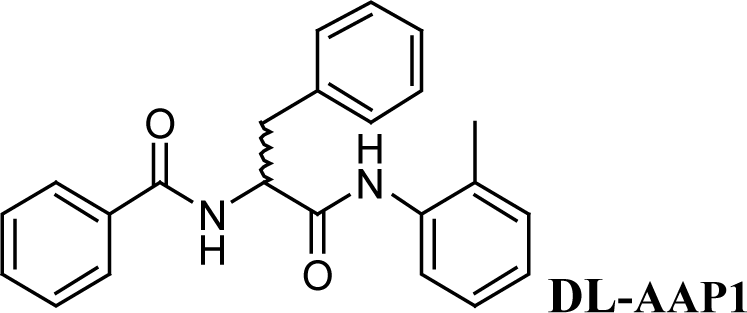

D/L-Benzoyl phenylalanine (32.2 mg; 120 μmol; Chem-Impex, Inc.) was dissolved in 2 ml anhydrous DMF at 25^o^C. To the solution, was added PS-carbodiimide (180 mg, 240 μmol; Biotage, Inc.) and hydroxybenzotriazole (24.3 mg; 180 μmol; Sigma-Aldrich, Inc.). The reaction mixture was stirred for 15 min under argon, o-toluidine (10.63 μ!, 100 μmol; Sigma-Aldrich, Inc.) was added, and stirring was continued for 2 h. The mixture was filtered through a plug of glass wool and evaporated to dryness. The sample was re-dissolved in chloroform and purified by silica chromatography with chloroform as eluent. Yield: 31.9 mg, 89%. ^1^H-NMR (500 MHz, CDCl3): δ 2.18 (s, 3H), 3.25 (dd, 1H), 3.35 (dd, 1H), 5.05 (dd, 1H), 6.8-7.8 (m; 14H, aryl protons). MS (MALDI): calculated: *m*/*z* 359.43 (Mtf); found: 359.35, 381.33 (M+Na^+^).

#### Nα-benzoyl-N-(2-methylphenyl)-D-phenylalaninamide (D-AAP1)

**D-AAP1.**
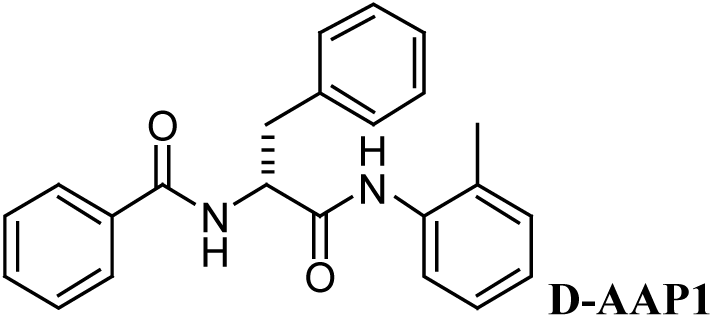

D-AAP1 was prepared as for DL-AAP1, but using D-benzoyl phenylalanine (Chem-Impex, Inc.) in place of D/L-benzoyl phenylalanine. The product (100 μg) was dissolved in isopropanol (100 μl and purified by chiral HPLC on a 4.6 x 250 mm ChiralPak IA column (ChiralPak, Inc.) in 10% isopropanol in hexanes at 1 ml/min. Two peaks were observed, one exhibiting a retention time of 14 min and an integrated area of 1.1 arbitrary units (assigned as the D stereoisomer, D-AAP1; 10% enantiomeric excess), and another exhibiting a retention time of 17 min and an integrated area of 1.0 arbitrary unit (assigned as the L stereoisomer, L-AAP1). The peak assigned as the D stereoisomer was collected. MS (MALDI): calculated: *m*/*z* 359.43 (MH^+^); found: 359.35, 381.33 (M+Na^+^). Optical rotation in dimethylformamide, 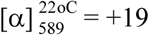.

#### Nα-benzoyl-N-(2-methylphenyl)-L-phenylalaninamide (L-AAP1)

**L-AAP1.**
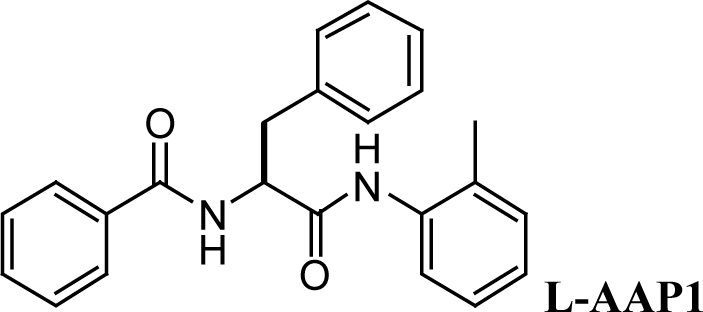

L-AAP1 was prepared as for D-AAP1, but using L-benzoyl phenylalanine (Chem-Impex, Inc.) in place of D-benzoyl phenylalanine. Two peaks were observed, one exhibiting a retention time of 14 min and an integrated area of 1.0 arbitrary unit (assigned as the D stereoisomer, D-AAP1), and another exhibiting a retention time of 17 min and an integrated area of 1.1 arbitrary units (assigned as the L stereoisomer, L-AAP1; 10% enantiomeric excess). The peak assigned as the L stereoisomer was collected. MS (MALDI): calculated: *m*/*z* 359.43 (MH^+^ found: 359.35, 381.33 (M+Na^+^). Optical rotation in dimethylformamide, 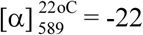.

#### Nα-selenophenoyl-N-(2-methylphenyl)-D-phenylalaninamide (D-IX336)

**D-IX336.**
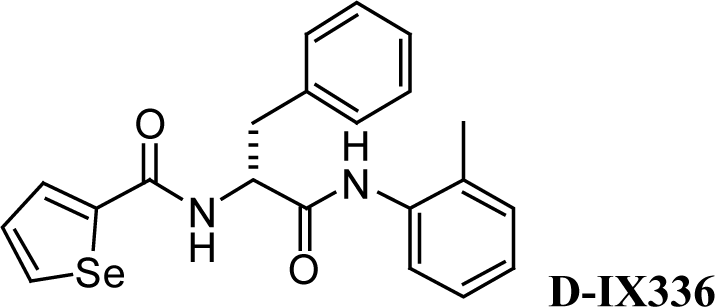

To a suspension of N-Fmoc-D-phenylalanine (2 g, 5.16 mmol; Chem-Impex, Inc.) in 20 ml dichloromethane was added oxalyl chloride (0.66 ml, 7.74 mmol; Sigma-Aldrich, Inc.) and 200 μ! dimethylformamide. After stirring for 30 min, the reaction mixture was evaporated to dryness (to remove any trace of unreacted oxalyl chloride) and re-dissolved in 30 ml dichloromethane. O-toluidine (0.554 ml, 5.16 mm; Sigma-Aldrich, Inc.) was added drop-wise followed by addition of DIPEA (1.08 ml, 6.19 mm). The reaction was stirred for 2 h, and 0.5 M HCl was used to acidify the mixture, which was then extracted with 3 x 30 ml dichloromethane. The organic extract was filtered, washed with brine, dried with anhydrous sodium sulfate, and evaporated to a solid, providing (9H-fluoren-9-yl) methyl (R)-(1-oxo-3-phenyl-1-(o-tolylamino) propan-2-yl) carbamate, which was used without purification in the next step. Crude yield: 1.2 g, 49%. MS (MALDI): calculated: *m*/*z* 476.58 (MH^+^); found: 477.13.

To 0.2 g (9H-fluoren-9-yl) methyl (R)-(1-oxo-3-phenyl-1-(o-tolylamino)-propan-2-yl) carbamate in 15 ml dichloromethane was added 0.16 ml piperidine. The reaction mixture was stirred for 6 h, evaporated, and purified by silica chromatography with 2% methanol in dichloromethane. The lower-eluting fractions were pooled and re-purified by silica chromatography (25-100% ethyl acetate/hexanes), providing (R)-2-amino-3-phenyl-N-(o-tolyl)propanamide. Yield: 17.6 mg; 17%. mS (MALDI): calculated: *m*/*z* 412.36 (MH^+^); found: 413.01.06, 434.99 (M+Na^+^).

To (R)-2-amino-3-phenyl-N-(o-tolyl)-propanamide (17.6 mg, 69 μmol) in 0.5 ml dichloromethane, was added selenophene-2-carboxylic acid (18.2 mg in 0.5 mL dichloromethane, 104 μmol, LabNetwork), DIPEA (18.2 μλ, 104 μmol) and propane phosphonic acid anhydride (T3P; 88.6 μλ, 140 μmol; Sigma-Aldrich, Inc. as 50% solution in ethyl acetate). The reaction mixture was stirred for 22 h and then evaporated to an oil. The resulting oil was re-dissolved in 10 ml ethyl acetate, washed with 5 ml 0.5M HCl, 5 ml saturated sodium bicarbonate, 5 ml brine, dried with anhydrous sodium sulfate, and evaporated to dryness. The crude material was purified by semi-preparative reverse-phase HPLC (25 x 10mm Jupiter C18, 10 μμ, 300 Å; 30-60% B in 30 min; A = 0.1% trifluoroacetic acid; B = 0.1% trifluoroacetic acid in acetonitrile; 4 ml/min; Phenomenex, Inc.), providing D-IX336. Yield: 7 mg; 25%. MS (MALDI): calculated: *m*/*z* 254.33 (MH^+^); found: 255.06, 277.03 (M+Na^+^). ^1^H-NMR (500 MHz, CDCl_3_): δ 8.22 (d, 1H), 7.79 (d, 1H), 7.65 (s, 1H), 7.55(br s, 1H), 7.00-7.40 (m, 9H), 6.70 (d, 1H), 4.80 (m, 1H), 3.20 (dd, 1H), 3.38 (dd, 1H), 1.99 (s, 3H).

##### *M. tuberculosis* σ^A^

*E. coli* strain BL21(DE3) (Invitrogen, Inc.) was transformed with plasmid pET30a-*Mtb*-σ^A^ (*19*), encoding N-terminally hexahistidine-tagged*Mtb* σ^A^ under control of the bacteriophage T7 gene 10 promoter. Single colonies of the resulting transformants were used to inoculate 50 ml LB broth containing 50 μg/ml kanamycin, and cultures were incubated 16 h at 37°C with shaking. Aliquots (10 ml) were used to inoculate 1 L LB broth containing 50 μg/ml kanamycin, cultures were incubated at 37°C with shaking until OD_600_ = 0.8, cultures were induced by addition of IPTG to 1 mM, and cultures were incubated overnight at 16°C. Cells were harvested by centrifugation (4,000 x g; 15 min at 4 °C), re-suspended in 20 ml buffer A (10 mM Tris-HCl, pH 7.9, 0.2 M NaCl, 5% glycerol), and lysed using an EmulsiFlex-C5 cell disrupter (Avestin, Inc.). The lysate was centrifuged (20,000 x g; 30 min at 4°C), and the supernatant was loaded onto a 5 ml column of Ni-NTA-agarose (Qiagen, Inc.) equilibrated in buffer A. The column was washed with 50 ml buffer A containing 40 mM imidazole and eluted with 25 ml buffer A containing 0.25 M imidazole. The sample was further purified by anion-exchange chromatography on a 16/10 Mono Q column (GE Healthcare, Inc.; 160 ml linear gradient of 300-500 mM NaCl in 10 mM Tris-HCl, pH 7.9, 0.1 mM EDTA, 5 mM dithiothreitol, and 5% glycerol; flow rate = 1 ml/min). Fractions containing σ^A^ were pooled, were concentrated to ~15 mg/ml using 30 kDa MWCO Amicon Ultra-15 centrifugal ultrafilters (Millipore, Inc.) and stored at −80°C. Yields were ~10 mg/L, and purities were >95%.

*Mtb* σ^A^ was prepared analogously, using plasmid pET30a-*Mtb*-σ^A^ΔR1.1), encoding N-terminally hexahistidine-tagged *Mtb* σ^A^(225-528) under control of the bacteriophage T7 gene 10 promoter [constructed by replacing the *Eco*RI-*Not*I segment of plasmid pET30a (EMD Millipore, Inc.) by the *Eco*RI-*Not*I DNA segment of a DNA fragment carrying GAATTC, followed by codons 225-528 of *Mtb sigA*, followed by TAAGCGGCCGC, generated by PCR with plasmid pET30a-*Mtb*-σ^A^ as template].

##### *M. tuberculosis* β’*Mtb*SI

*E. coli* strain BL21 (DE3) (Invitrogen, Inc.) was transformed with plasmid pET28a-Mtb-β’MtbSI, encoding N-hexahistidine-tagged β’MtbSI under control of the bacteriophage T7 gene 10 promoter [constructed by replacing the *Nde*I-*Eco*R segment of plasmid pET28a (EMD Millipore, Inc.) with the *Nde*I-*Eco*RI DNA segment of a DNA fragment carrying CATATG, followed by codons 141-229 of *Mtb rpoC*, followed by TAAGAATTC, generated by PCR with plasmid pCOLADuet-*rpoBC* (*19*) as template]. Single colonies of the resulting transformants were used to inoculate 50 ml LB broth containing 50 μg/ml kanamycin, and cultures were incubated 16 h at 37°C with shaking. Aliquots (10 ml) were used to inoculate 1 L LB broth containing 50 μg/ml kanamycin, cultures were incubated at 37°C with shaking until OD_600_ = 0.8, cultures were induced by addition of IPTG to 1 mM, and cultures were incubated overnight at 16°C. Cells were harvested by centrifugation (4,000 x g; 15 min at 4°C), resuspended in 20 ml buffer A (10 mM Tris-HCl, pH 7.9, 0.2 M NaCl, 5% glycerol), and lysed using an EmulsiFlex-C5 cell disrupter (Avestin, Inc.). The lysate was centrifuged (20,000 x g; 30 min at 4°C), and the supernatant was loaded onto a 5 ml column of Ni-NTA-agarose (Qiagen, Inc.) equilibrated in buffer A. The column was washed with 50 ml buffer A containing 40 mM imidazole and eluted with 25 ml buffer A containing 0.25 M imidazole. The sample was further purified using by gel filtration on a Superdex G500 column (GE Healthcare, Inc) in 20 mM Tris-HCl, pH 8.0, 75 mM NaCl, 5 mM MgCl_2_, and was concentrated to ~10 mg/L in the same buffer using 10 kDa MWCO Amicon Ultra-15 centrifugal ultrafilters (Millipore, Inc.) and stored at −80°C. Yields were ~20 mg/L, and purities were >95%.

Selenomethionine-substituted β’*Mtb*SI protein was produced using plasmid pET28a-Mtb-β’*MtbS* and the procedures of *20* and was purified as for native β’*Mtb*SI.

##### *M. tuberculosis* RNAP core enzyme

*Mtb* RNAP core enzyme and RNAP core enzyme derivatives were prepared from cultures of *E. coli* strain BL21 (DE3) (Invitrogen, Inc.) co-transformed with plasmids pACYC-<u>σ^A^</u>, pCOLADuet-*rpoBC*, and pCDF-*rpoZ* (*21*), or derivatives thereof constructed using site-directed mutagenesis (QuickChange Kit; Agilent Biotechnology, Inc.), using procedures as in *21*.

##### *M. tuberculosis* RNAP σ^A^ holoenzyme

*Mtb* RNAP σ^A^ holoenzyme and RNAP σ^A^ holoenzyme derivatives were prepared from cultures of *E. coli* strain BL21 (DE3) (Invitrogen, Inc.) co-transformed with plasmids pACYC-*rpoA-<u>σ^A^</u>*, pETDuet-*rpoBC* and pCDF-*rpoZ* (*21*), or derivatives thereof constructed using site-directed mutagenesis (QuikChange Kit; Agilent Biotechnology, Inc.), using procedures as in *21*.

##### Oligonucleotides

Oligodeoxyribonucleotides (IDT, Inc.) and oligoribonucleotides (GpA, GpGpA, and UpCpGpA; Trilink, Inc.) were dissolved in nuclease-free water (Ambion, Inc.) to 3 mM and stored at −80°C.

##### Nucleic-acid scaffolds

Nucleic-acid scaffolds RPo, RPitc3, and RPitc4 (sequences in Fig. S1C) were prepared as follows: Nontemplate-strand oligodeoxyribonucleotide (0.5 mM), template-strand oligodeoxyribonucleotide (0.55 mM), and, where indicated, 3 nt or 4 nt oligoribonucleotide (1 mM) in 40 μl 5 mM Tris-HCl, pH 7.7, 0.2 M NaCl, and 10 mM MgCl_2_ were heated 5 min at 95°C, cooled to 25°C in 2°C steps with 1 min per step using a thermal cycler (Applied Biosystems, Inc.), and stored at −80°C.

##### RNAP-DNA interaction assays

Stabilities of *Mtb* RPo were assessed using fluorescence polarization assays (*22*) in a 96-well microplate format. Reaction mixtures contained (100 μl: 0-320 μM*Mtb* RNAP σ^A^ holoenzyme or derivative, 10 nM Cy3-labelled nucleic-acid scaffold RPo (sequence in Fig. S1C; Cy3 incorporated at 5′ end of nontemplate strand), 20 mM Tris-HCl (pH 8.0), 250 mM NaCl, 5 mM MgCl_2_, and 5 mM dithiothreitol. Reaction mixtures were incubated 10 min at 25°C, and fluorescence emission intensities were measured using a microplate reader equipped with polarizers (GENios Pro; TECAN, Inc; excitation wavelength = 550 nm; emission wavelength = 570 nm). Fluorescence polarization was calculated using:

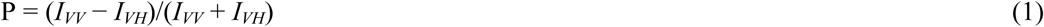

where *I_VV_* and *I_VH_* are fluorescence intensities with the excitation polarizer at the vertical position and the emission polarizer at, respectively, the vertical position and the horizontal position.

Equilibrium dissociation constants, K_D_, were extracted by non-linear regression using:

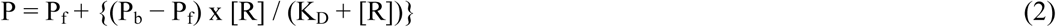

where P is fluorescence polarization at RNAP concentration R, P_f_ is fluorescence polarization in the absence of RNAP, and P_b_ is the fluorescence polarization in the presence of a saturating RNAP concentration.

##### RNAP-inhibitory activities

Fluorescence-detected RNAP-inhibition assays were performed using the profluorescent substrate γ-{2′-(2-benzothiazoyl)-60-hydroxybenzothiazole]-ATP [BBT-ATP; procedures as in *17*, using 150 nM *Mtb* RNAP core enzyme plus 600 nM*Mtb* σ^A^, 75 nM *Eco* RNAP σ^70^ holoenzyme (prepared as in *23*), or 75 nM *S. aureus* RNAP core enzyme and 300 nM *S. aureus* σ^A^ (prepared as in *24*) and 20 nM DNA fragment containing bacteriophage T5 N25 promoter (prepared as in *23*)].

Radiochemical assays with human RNAP I/II/III were performed as in *25*.

Half-maximal inhibitory concentrations (IC50s) were calculated by non-linear regression in SigmaPlot (SPSS, Chicago, IL).

##### Growth-inhibitory activities

Minimal inhibitory concentrations (MICs) for *Mycobacterium tuberculosis* H37Rv and*M. avium* ATCC 25291 were quantified in microplate Alamar Blue assays (procedures as in *26*).

MICs for *M. smegmatis* mc^2^ 155, *M. smegmatis* ATCC 19420, and derivatives thereof were quantified in broth microdilution assays [procedures essentially as in *27*, using 7H9 broth (BD Biosciences, Inc.) supplemented with 0.5% glycerol, and 0.05 % Tween-80, and using a starting cell density of 2-5×10^5^ cfu/ml, and incubation 72 h at 37°C.

MICs for *Staphylococcus aureus* ATCC 12600, *Escherichia coli* D21f2tolC, and mammalian cells (Vero E6) were determined as in *25*.

##### Resistance and cross-resistance assays

*M. smegmatis* mc^2^ 155 (*rpoB′637′A*), a derivative of*M. smegmatis* mc^2^ 155 having a chromosomal rpoB that encodes RNAP β subunit having a substitution corresponding to Eco β R637A, was constructed using recombineering with targeting oligonucleotide 5′-GGCGGAGACGAACTCGACCTCGCCGCCCTTCTTGGCGACCATGACGCGGTCCTCGGTGAAGCGGCCGTTCTC-3′ [procedures as in *28*; selected on Seven H11 agar (BD Biosciences, Inc.) containing Middlebrook ADC enrichment (BD Biosciences, Inc.), 0.5% glycerol, and 30 μg/ml D-AAP1; confirmed by PCR amplification and sequencing or *rpoB* and *rpoC*].

*M. smegmatis* ATCC 19420 (*rpoB′526′Y*) and ATCC 19420 (*rpoB′531′L*, ATCC 19420 derivatives having chromosomal *rpoB* genes that encode RNAP β subunits having substitutions that correspond to *Eco* β H526Y and *Eco* β S5331L, were obtained as spontaneous Rif-resistant mutants (selected on Seven H11 agar containing Middlebrook ADC enrichment, 0.5% glycerol, and 50 μg/ml Rif; confirmed by PCR amplification and sequencing of *rpoB* and rpoC].

Resistance and cross-resistance levels were determined in broth microdilution assays (see above, “Growth-inhibitory activities”), using final concentrations of 0.0015-50 μg/ml D-AAP or Rif.

##### Checkerboard interaction assays

To assess antibacterial activities of combinations of D-AAP1 and Rif, broth-microdilution assays with *M. smegmatis* ATCC 19420 (see above, “Growth inhibitory activities”) were performed in a checkerboard format (*29-31*), analyzing in quintuplicate, all pairwise combinations of D-AAP1 at 4.00x, 3.50x, 3.00x, 2.67x, 2.63x, 2.50x, 2.33x, 2.25x, 2.00x, 1.88x, 1.75x, 1.67x, 1.50x, 1.33x, 1.25x, 1.13x, 1.00x, 0.750x, 0.667x, 0.500x, 0.375x, 0.333x, and 0.250 MIC (MIC_D-AApP1_ = 0.20 μg/ml) with Rif at 1.33x, 1.17x, 1.00x, 0.875x, 0.833x, 0.750x, 0.667x, 0.656x, 0.625x, 0.500x, 0.469x, 0.375x, 0.333x, 0.281x, 0.250x, 0.188x, 0.167x, 0.125x, and 0.094x MIC (MIC_Rif_ = 3.13 μg/ml). Minimum and maximum fractional inhibitory concentrations indices (FICI_max_ and FICI_min_) were calculated as in *30*. FICI_min_ ≤ 0.5 was deemed indicative of super-additivity (synergism), FICI_min_ > 0.5 and FICI_max_ ≤ 4.0 was deemed indicative of additivity, and FICI_max_ > 4.0 was deemed indicative of sub-additivity (antagonism) (*30-31*).

##### Resistance-rate assays

Resistance rates were determined using fluctuation assays essentially as in *32*. Defined numbers of cells of *M. smegmatis* ATCC 19420 (~10^9^ cfu/plate) were plated on Seven H11 agar containing Middlebrook ADC enrichment, 0.5% glycerol, and either 1.56 μg/ml D-AAP1, 50 μg/ml Rif, or both 1.56 μg/ml D-AAP1 and 50 μg/ml Rif; and numbers of colonies were counted after 72 h at 37°C (at least 4 determinations for each concentration of each test compound). Resistance rates were calculated using the Ma-Sandri-Sarkar Maximum Likelihood Estimator (*33-34*) as implemented on the Fluctuation Analysis Calculator (FALCOR; www.keshavsingh.org/protocols/FALCOR.html; *35*). Sampling correction was performed as in *36*.

##### Structure determination: assembly of transcription initiation complexes

For *Mtb* RPo, RPitc3, and RPitc4, complexes for crystallization were prepared by mixing 16 μl 50 μM*Mtb* RNAP σ^A^ holoenzyme (in 20 mM Tris-HCl, pH 8.0, 75 mM NaCl, 5 mM MgCl_2_, and 5 mM dithiothreitol,) and 4 μl 0.4 mM of nucleic-acid scaffold RPo, RPitc3, or RPitc4 (in 5 mM Tris-HCl, pH 7.7, 0.2 M NaCl, and 10 mM MgCl_2_) and incubating 1 h at 25°C. For *Mtb* RPitc2, the complex for crystallization was prepared by mixing 16 μl 50 μM*M. tuberculosis* RNAP holoenzyme (in 20 mM Tris-HCl, pH 8.0, 75 mM NaCl, 5 mM dithiothreitol, 5 mM MgCl_2_), 4 μl 0.4 mM of nucleic-acid scaffold RPo (in 5 mM Tris-HCl, pH 7.7, 0.2 M NaCl, and 10 mM MgCl_2_), and 1 μl 25 mM GpA (in water), and incubating 1 h at 25°C. For *Mtb* RPo, RPitc3, and RPitc4 in complex with Rif, complexes for crystallization were prepared by incubating 16 μl 50 μM*Mtb* RNAP σ^A^ holoenzyme (in 20 mM Tris-HCl, pH 8.0, 75 mM NaCl, 5 mM MgCl_2_, and 5 mM dithiothreitol,) and 0.5 μl 8 mM Rif (Sigma-Aldrich, Inc.) for 0.5 h at 25°C, and then adding 4 μl 0.4 mM of nucleic-acid scaffold RPo, RPitc3, or RPitc4 (in 5 mM Tris-HCl, pH 7.7, 0.2 M NaCl, and 10 mM MgCl_2_), and incubating 1 h at 25°C. For *Mtb* RPitc,2 in complex with Rif, the complex for crystallization was prepared by incubating 16 μl 50 μM*Mtb* RNAP σ^A^ holoenzyme (in 20 mM Tris-HCl, pH 8.0, 75 mM NaCl, 5 mM MgCl_2_, and 5 mM dithiothreitol) and 0.5 μl 8 mM Rif for 0.5 h at 25°C, and then adding 4 μl 0.4 mM of nucleic-acid scaffold RPo (in 5 mM Tris-HCl, pH 7.7, 0.2 M NaCl, and 10 mM MgCl_2_) and 1 μl 25 mM GpA (in water) and incubating 1 h at 25 °C.

##### Structure determination: crystallization, cryo-cooling, and crystal soaking

Robotic crystallization trials were performed for *Mtb* RPo and *Mtb* β’MtbSI using a Gryphon liquid handling system (Art Robbins Instruments, Inc.), commercial screening solutions (Emerald Biosystems, Inc.; Hampton Research, Inc.; and Qiagen, Inc.), and the sitting-drop vapor diffusion technique [drop: 0.2 μl transcription initiation complex (previous section) or 0.2 μl 500 μM *Mtb* β’*Mtb*SI (in 20 mM Tris-HCl, pH 8.0, 75 mM NaCl, 5 mM MgCl_2_, and 5 mM dithiothreitol) plus 0.2 μl screening solution; reservoir: 60 μl screening solution; 22°C]. 900 conditions were screened. Under several conditions, *Mtb* RPo crystals appeared within two week, and *Mtb* β’*Mtb*SI crystals appeared within one week. Conditions were optimized using the hanging-drop vapor-diffusion technique at 22°C. The optimized conditions for *Mtb* RPo (drop: 1 μl RPo in 20 mM Tris-HCl, pH 8.0, 75 mM NaCl, 5 mM MgCl_2_, and 5 mM dithiothreitol plus 1 μl 50 mM Tris-HCl, pH 8.2, 200 mM KCl, 50 mM MgCl_2_, and 10% PEG3350 plus 0.2 μ 80.0 mM CHAPSO; reservoir: 400 μ 50 mM Tris-HCl, pH 8.2, 200 mM KCl, 50 mM MgCl_2_, and 10% PEG3350; 22°C) yielded high-quality, rod-like crystals with dimensions of 0.4 mm x 0.1 mm x 0.1 mm in two weeks (Fig. S1C). The optimized conditions for β’*Mtb*SI (drop: 1 μl β’*Mtb*SI in 20 mM Tris-HCl, pH 8.0, 75 mM NaCl, 5 mM MgCl_2_ and 5 mM dithiothreitol plus 1 μl 100 mM sodium citrate tribasic dihydrate, pH 5.5, and 22% PEG1000; reservoir: 400 μl 100 mM sodium citrate tribasic dihydrate, pH 5.5, and 22% PEG1000; 22°C). *Mtb* RPo and *Mtb* β’*Mtb*SI crystals were transferred to reservoir solution containing 18% (v/v) (2R,3R)-(-)-2,3-butanediol (Sigma-Aldrich, Inc.) and flash-cooled with liquid nitrogen. Analogous procedures were used for *Mtb* RPitc2, RPitc3, RPitc4, and complexes with Rif.

For *Mtb* RPo in complex with D-AAP1 or D-IX336, *Mtb* RPo crystals were soaked by adding 0.2 μl 20 mM D-AAP1 or D-IX336 in (2R,3R)-(-)-2, 3-butanediol directly into the drop for 2 h, transferring to reservoir solution containing 18% (v/v) (2R,3R)-(-)-2,3-butanediol, and flash-cooling with liquid nitrogen. For *Mtb* RPo in complex with both Rif and D-AAP1, pre-formed crystals of *Mtb* RPo-Rif were soaked by adding 0.2 μl 20 mM of D-AAP1 in (2R,3R)-(-)-2,3-butanediol directly into the drop for 2 h, transferring to reservoir solution containing 18% (v/v) (2R,3R)-(-)-2,3-butanediol, and flash-cooling with liquid nitrogen.

##### Structure determination: data collection and reduction

Diffraction data and selenium single-wavelength anomalous dispersion data were collected from cryo-cooled crystals at Argonne National Laboratory beamline 19ID-D. Data were processed using HKL2000 (*37*). The resolution cutoff criteria were I/σ > 1.4 and R_merge_ < 1.

##### Structure determination: structure solution and refinement

The structure of *Mtb* RPo was solved by molecular replacement with Molrep (*38*) using the structure of *T. thermophilus* RPo (PDB 4G7H; 7) as the search model. One RNAP molecule was present in the asymmetric unit. Early-stage refinement included rigid-body refinement of RNAP, followed by rigid-body refinement of RNAP subunits, followed by rigid-body refinement of 44 RNAP domains (methods as in *7*). Electron density for nucleic acids was unambiguous, but was not included in models in early-stage refinement. Cycles of iterative model building with Coot (*39*) and refinement with Phenix (*40*) were performed. Improvement of the coordinate model resulted in improvement of phasing, and electron density maps for nucleic acids, which were not included in models at this stage of model building and refinement, improved over successive cycles. Nucleic acids then were built into the model and refined in stepwise fashion. The final model was generated by XYZ-coordinate refinement with secondary-structure restraints, followed by group B-factor and individual B-factor refinement. The final model, refined to R_work_ and R_free_ of 0.23 and 0.28, respectively, was deposited in the PDB with accession code (Table S1). Analogous procedures were used to solve and refine structures of *Mtb* RPitc2, RPitc3, and RPitc4 (Table S1). Analogous procedures were used to solve and preliminarily refine structures of complexes with inhibitors; models of inhibitors then were built into mF_o_-DF_c_ difference maps, and additional cycles of refinement and model building were performed, (Tables S2-S3).

The structure of β’*Mtb*SI was solved by the single-wavelength anomalous dispersion method, using Autosol as implemented in Phenix (*40*). The structure model was built using the program Coot (*39*), and structure refinement was carried out using Phenix (*40*). The final model was refined to 2.2 Å resolution (Table S3).

## Supplementary Figures

**Fig. S1.**
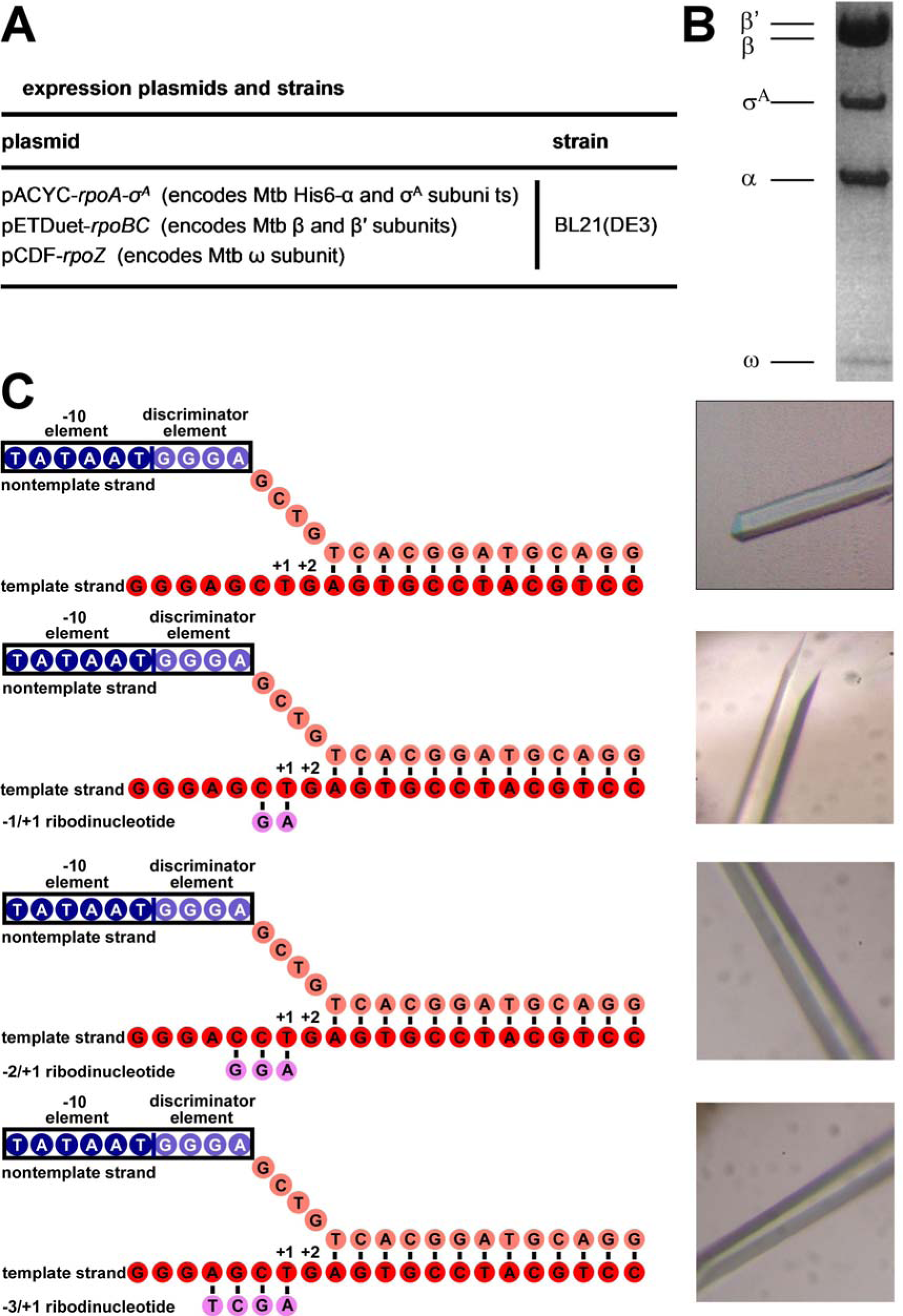
Structure determination: *Mtb* RPo, *Mtb* RPitc2, *Mtb* RPitc3, and *Mtb* RPitc4. (**A**) Plasmids and strain for production of *Mtb* RNAP σ^A^ holoenzyme in *E. coli* (*19,21*). (**B**) Coomassie-stained SDS-polyacrylamide gel electrophoresis of *Mtb* RNAP σ^A^ holoenzyme produced in *E. coli.* (**C**) Nucleic-acid scaffolds (left panels) and crystals (right panels).

**Fig. S2.**
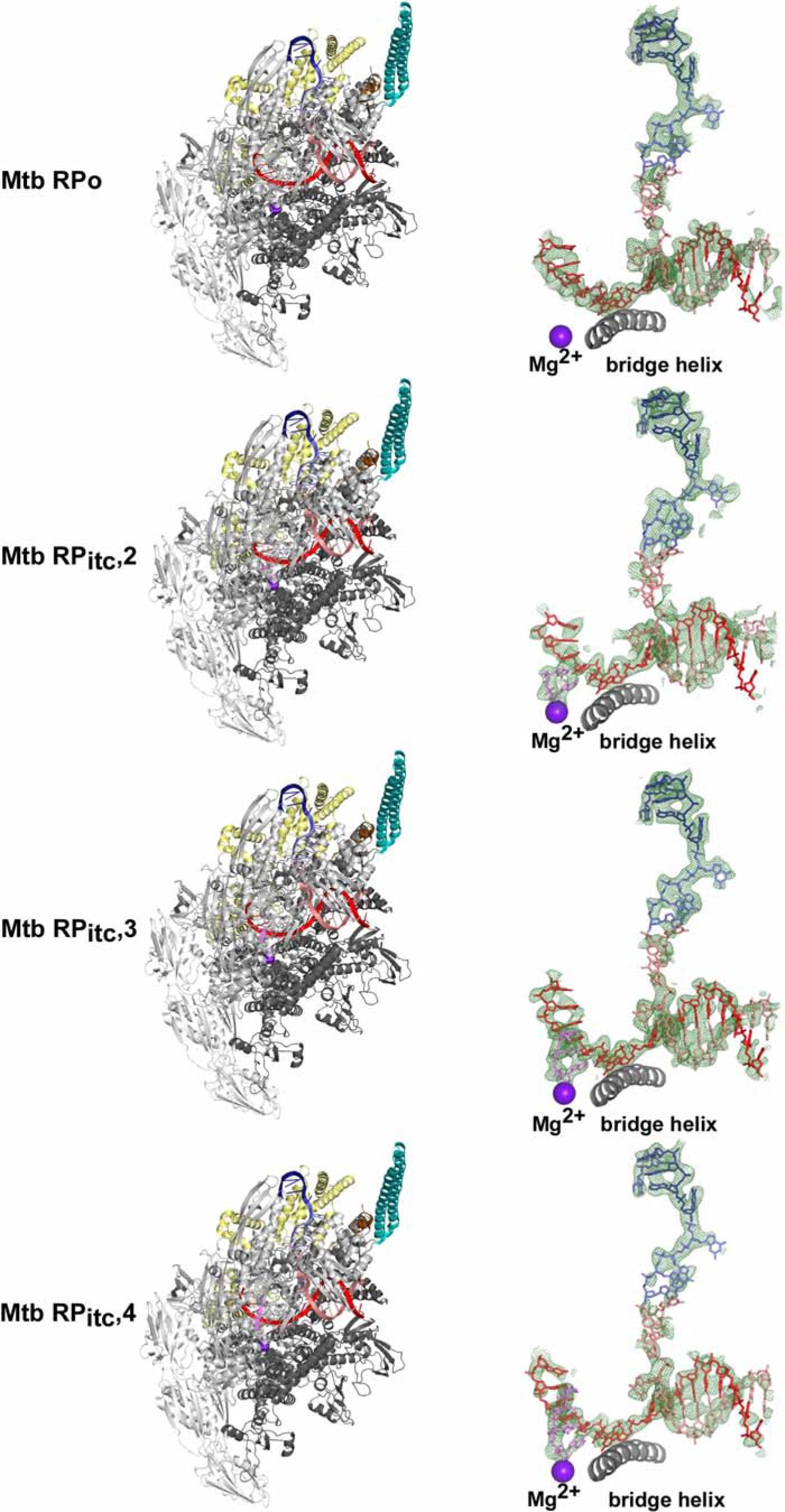
Structure determination: overall structures and electron density for *Mtb* RPo, RPitc2, RPitc3, and RPitc4. Left subpanels, overall structures. Colors as in Fig.1A. Right subpanels, electron density and model for nucleic acids.

**Fig. S3.**
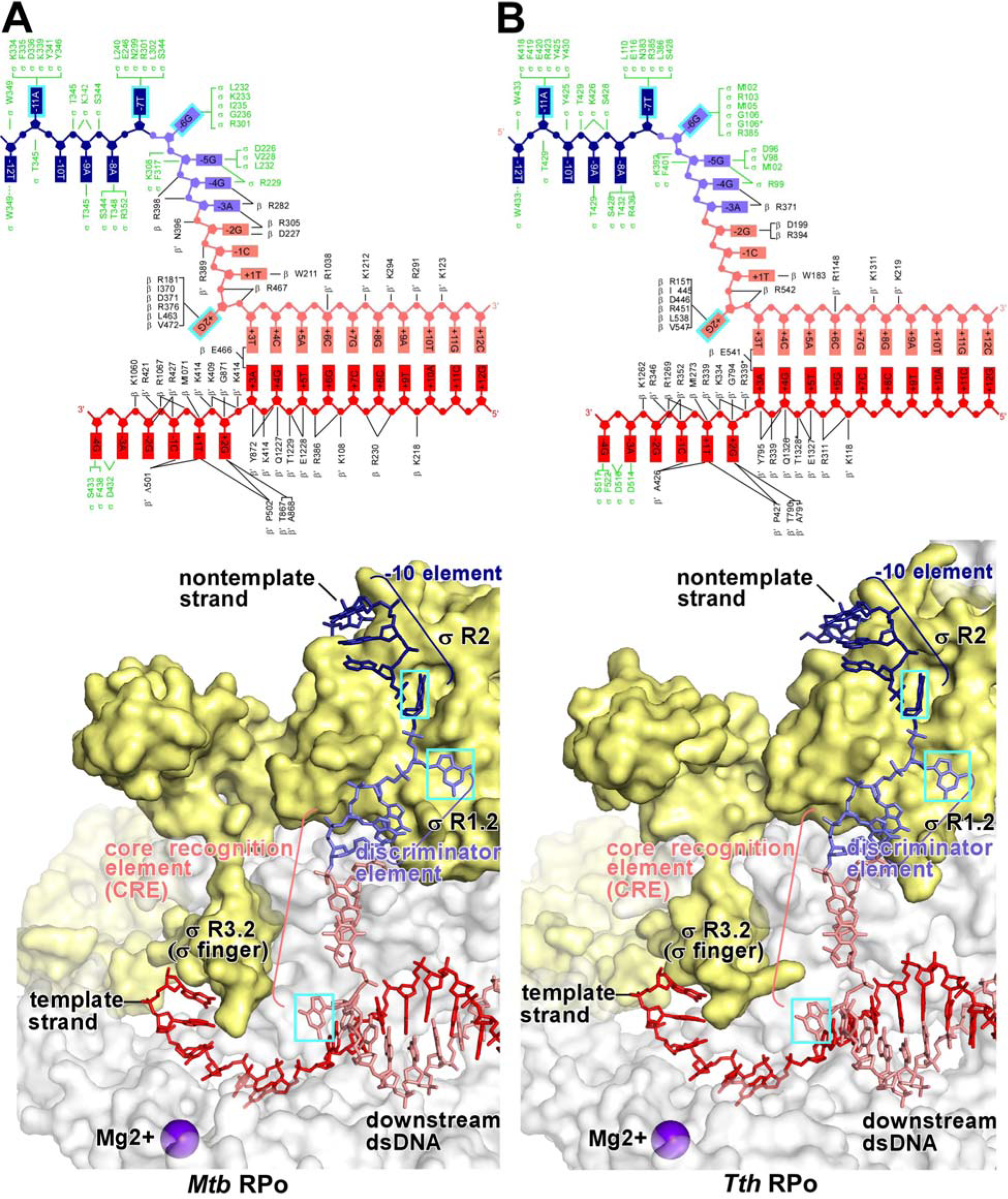
Comparison of structures of *Mtb* RPo and *Tth* RPo. (A) Top subpanel, summary of protein-nucleic-acid interactions in *Mtb* RPo. Black residue numbers and lines, interactions by *Mtb* RNAP; green residue numbers and lines, interactions by *Mtb* σ^A^; blue, −10 element of DNA nontemplate strand; light blue, discriminator element of DNA nontemplate strand; pink, rest of DNA nontemplate strand; red, DNA template strand; violet, active-center Mg^2+^; cyan boxes, bases unstacked and inserted into pockets. Residues numbered as in *Mtb* RNAP. Bottom subpanel, interactions of *Mtb* RNAP and *Mtb* σ^A^ with transcription-bubble non-template strand, transcription-bubble template strand, and downstream dsDNA. (B) Top subpanel, summary of protein-nucleic-acid interactions in *Tth* RPo (*7*). Residues numbered as in *Eco* RNAP. Bottom subpanel, interactions of *Tth* RNAP and Tth σ^A^ with transcription-bubble nontemplate strand, transcription-bubble template strand, and downstream dsDNA. Colors as in (A).

**Fig. S4.**
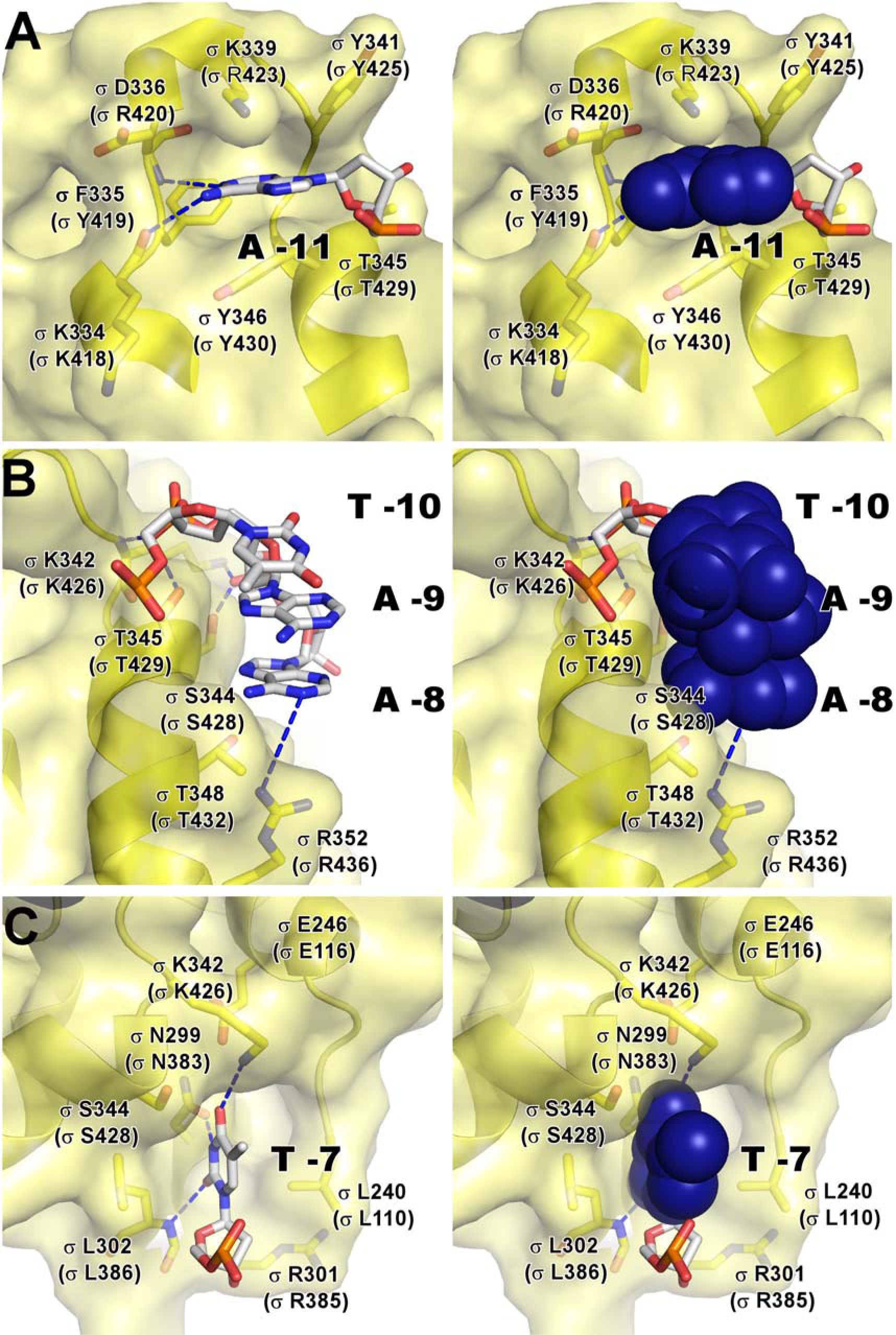
Recognition by *Mtb* σ^A^ of the −10 element: interactions with nontemplate-strand positions −11 through −7. In left subpanels, DNA nucleotides are shown in stick representation to highlight individual protein-nucleotide interactions. In right subpanels, DNA nucleotide base moieties are shown in space-filling representations to highlight protein-base steric complementarity. Yellow surfaces, solvent-accessible surfaces of *Mtb* σ^A^; dark blue surfaces, van der Waals surfaces of DNA bases; yellow ribbons, *Mtb* σ^A^ backbone; yellow, yellow-blue, and yellow-red stick representations, σ^A^ carbon, nitrogen, and oxygen atoms; white, blue, red, and orange stick representations, DNA carbon, nitrogen, oxygen, and phosphorous atoms; blue dashed lines, H-bonds. Coordinates are from the structure of *Mtb* RPo. (**A**) Interactions with nontemplate-strand position −11. (**B**) Interactions with nontemplate-strand positions −10, −9, and −8. (**C**) Interactions with nontemplate-strand positions −7.

**Fig. S5.**
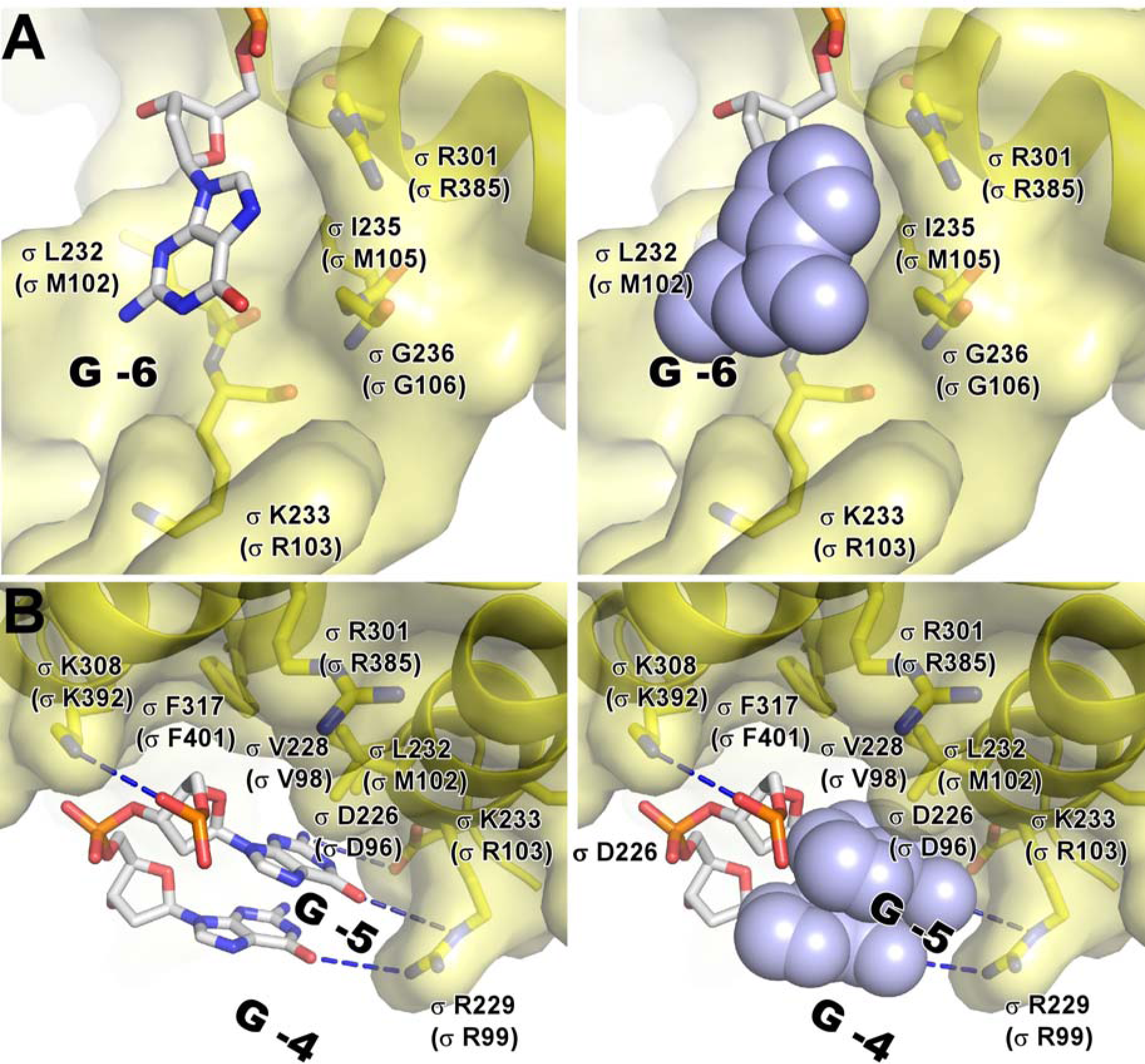
Recognition by *Mtb* σ^A^ of the discriminator element: interactions with nontemplate-strand positions −6 through −4. Subpanels and colors are as in Fig S4. Coordinates are from the structure of *Mtb* RPo. (**A**) Interactions with nontemplate-strand position −6. (**B**) Interactions with nontemplate-strand positions −5 and −4.

**Fig. S6.**
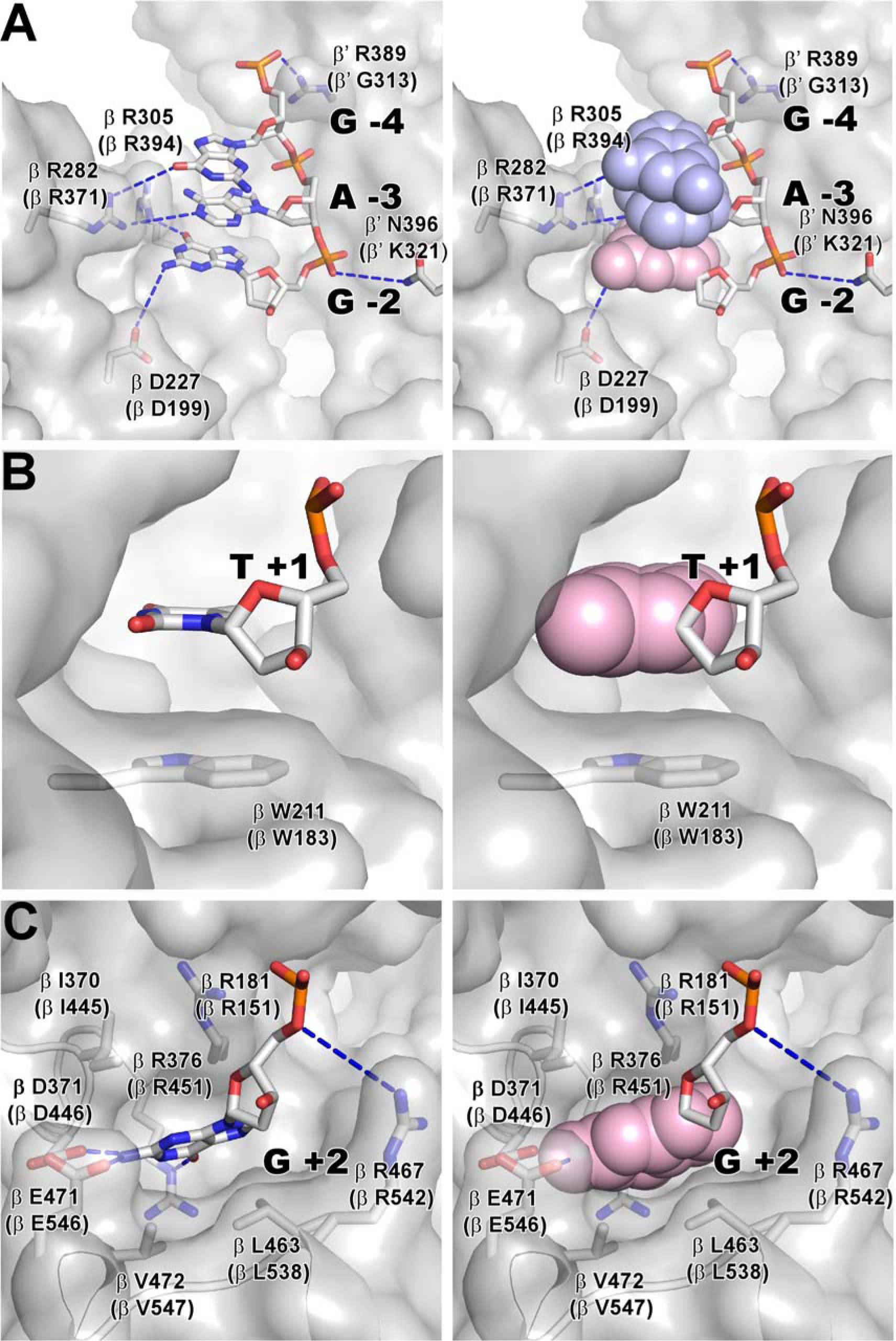
Recognition by *Mtb* σ^A^ of the core recognition element (CRE): interactions with nontemplate-strand positions −4 through +2. Subpanels and colors are as in Fig S4. Gray surfaces, solvent-accessible surfaces of RNAP β subunit; light blue and pink surfaces, van der Waals surfaces of DNA bases; gray ribbons, RNAP β-subunit backbone; gray, gray-blue, and gray-red stick representations, RNAP β-subunit carbon, nitrogen, and oxygen atoms. Other colors are as in Fig S4. Coordinates are from the structure of Mtb RPo. (**A**) Interactions with nontemplate-strand positions −4, −3 and −2. (**B**) Interactions with nontemplate-strand position +1. (**C**) Interactions with nontemplate-strand position +2.

**Fig. S7.**
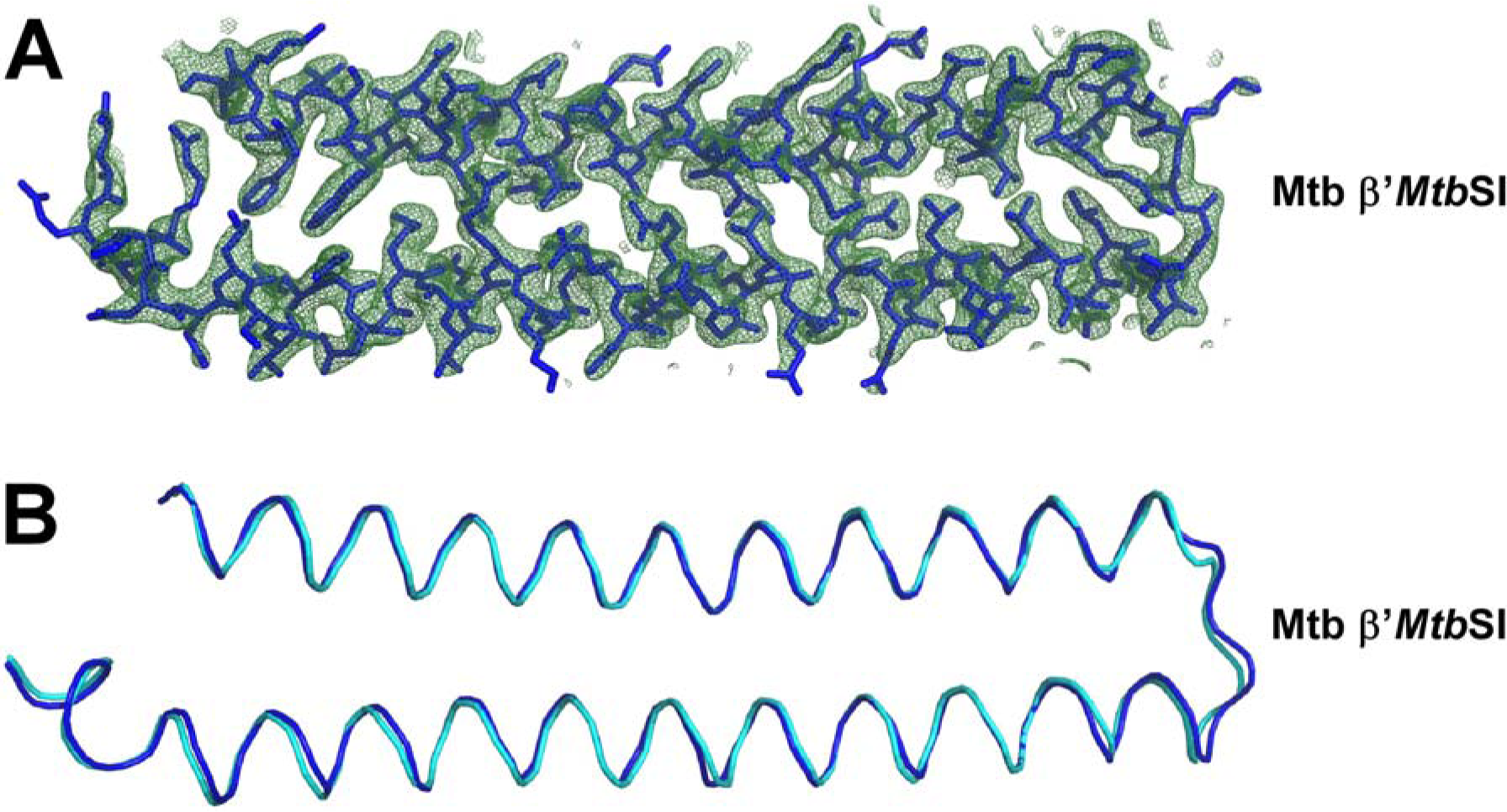
Structure of taxon-specific sequence insertion β′*Mtb*SI. (**A**) Electron density and model for β′*Mtb*SI from crystal structure of protein fragment *Mtb* β′(141-229). Blue sticks, β′*Mtb*SI atoms; green mesh, mF_o_-F_c_ electron density omit map (contoured at 2.5σ). (**B**) Superimposition of β′MtbSI from structure of *Mtb* RPo (cyan) on β′*Mtb*SI from structure of *Mtb* β′(141-229) (blue) (88 superimposed residues; rmsd = 0.65Â).

**Fig. S8.**
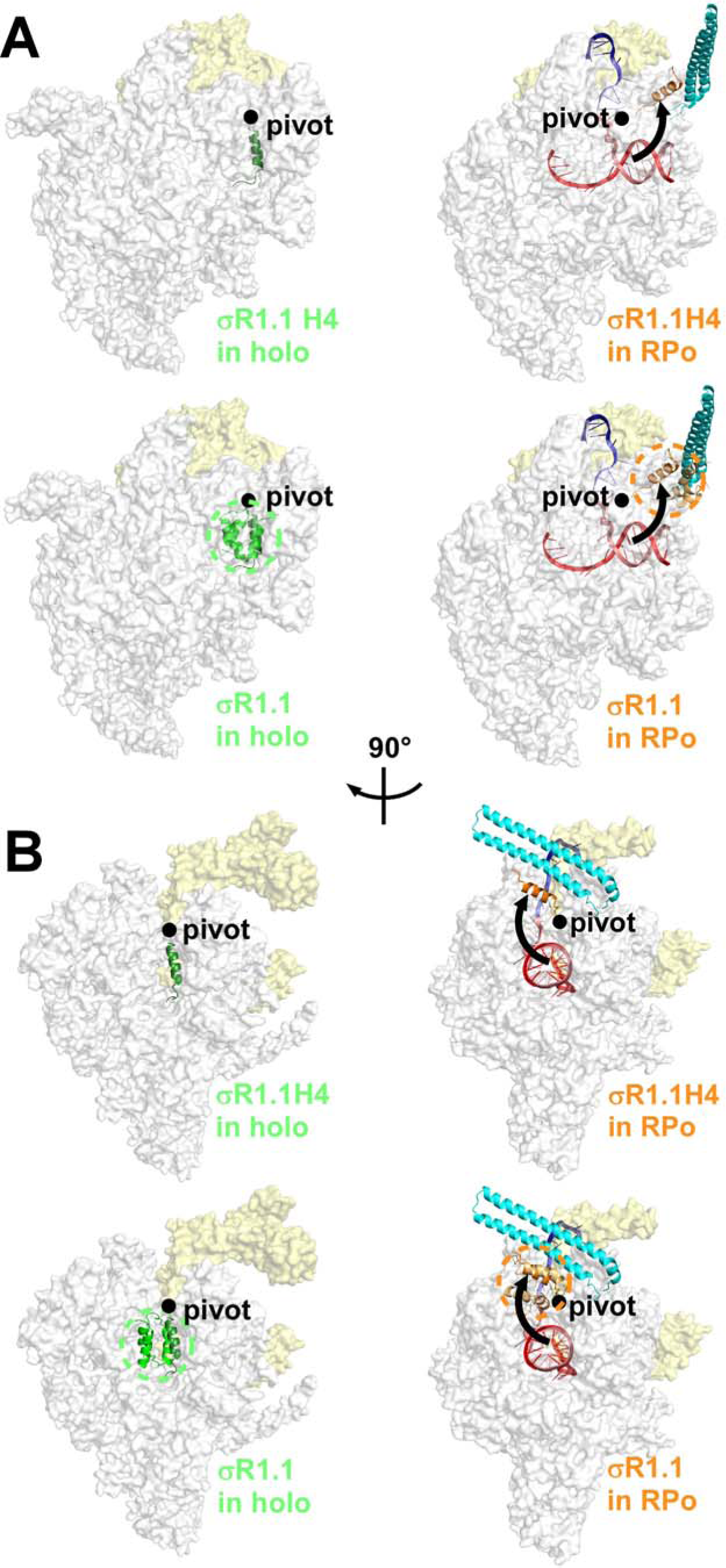
Difference in positions of σR1.1 in RNAP holoenzyme and RPo. (**A**) Top subpanels, structures of *Eco* RNAP holoenzyme (left) and *Mtb* RPo (right), showing position of σR1.1 helix 4 (H4; green in *Eco* RNAP holoenzyme and orange in *Mtb* RPo). Bottom subpanels, same as top subpanels, but showing full structure and approximate molecular volume of σR1.1 of *Eco* RNAP holoenzyme (green ribbons and green dashed circle) and H4 and approximate molecular volume of σR1.1 of *Mtb* RPo (orange ribbon and orange dashed circle). The position of σR1.1 H4 in RPo differs from the position in RNAP holoenzyme by a ~100° rotation (black arrow) about a “pivot” comprising the short unstructured segment between σR1.1 H4 and the rest of σ (black circle). (**B**) Same as (A), but orthogonal view.

**Fig. S9.**
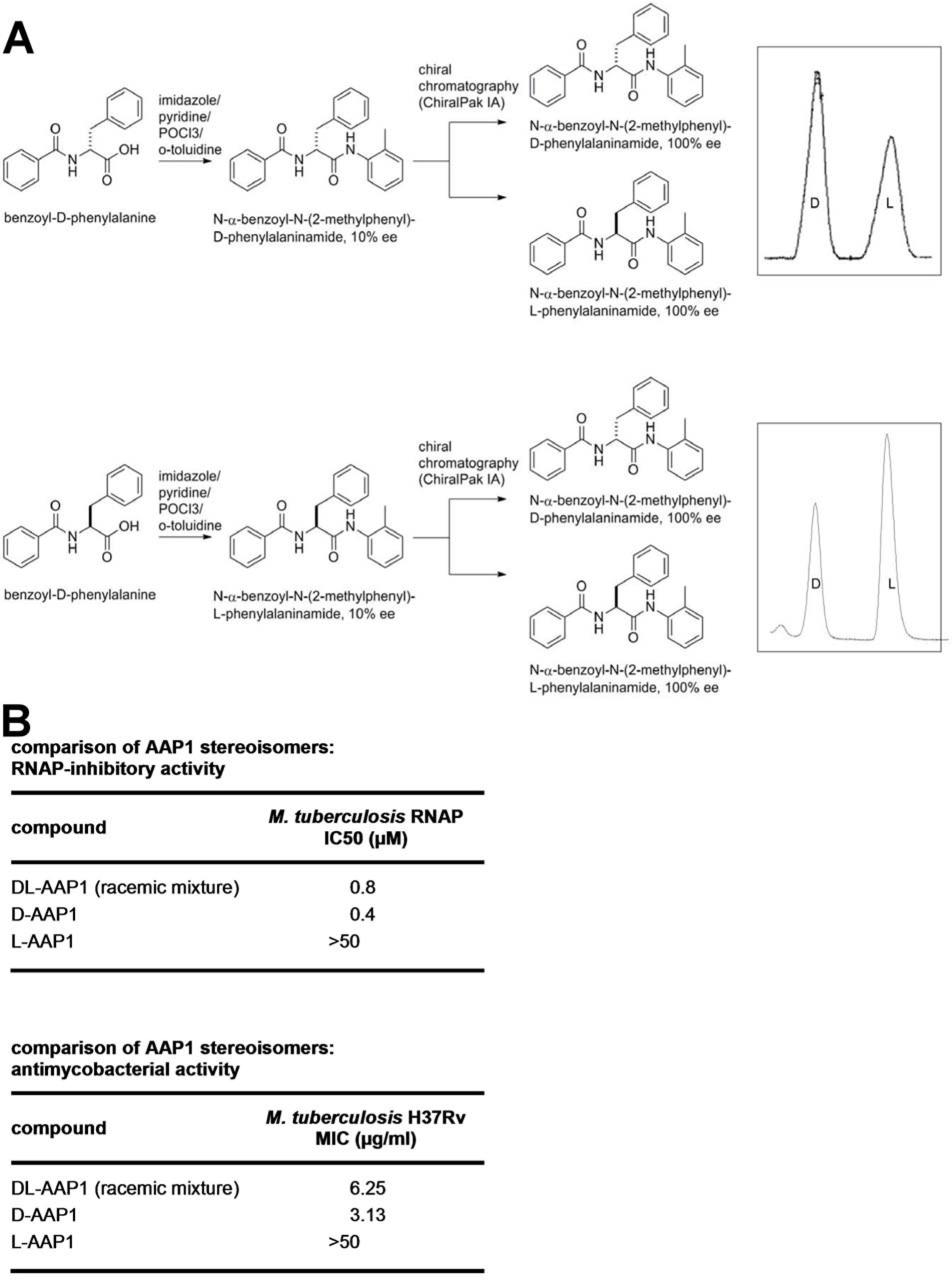
Preparation and stereospecificity of D-AAP1. (**A**) Upper panel, synthesis, chiral chromatography, and spectropolarimetry of D-AAP1 and L-AAP1 starting from D-stereoisomer precursor benzoyl-D-phenyalanine. Lower panel, synthesis an, chiral chromatography of D-AAP1 and L-AAP1 starting from L-stereoisomer precursor benzoyl-L-phenyalanine (**B**) RNAP-inhibitory activities and antibacterial activities of DL-AAP1, D-AAP1, and L-AAP1.

**Fig. S10.**
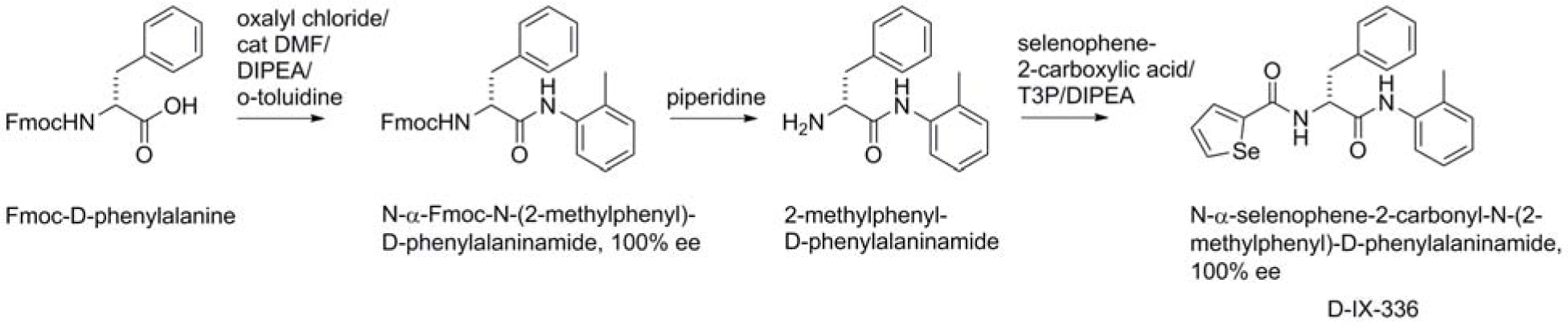
Preparation of selenium-containing D-AAP1 analog D-IX336.

**Fig. S11.**
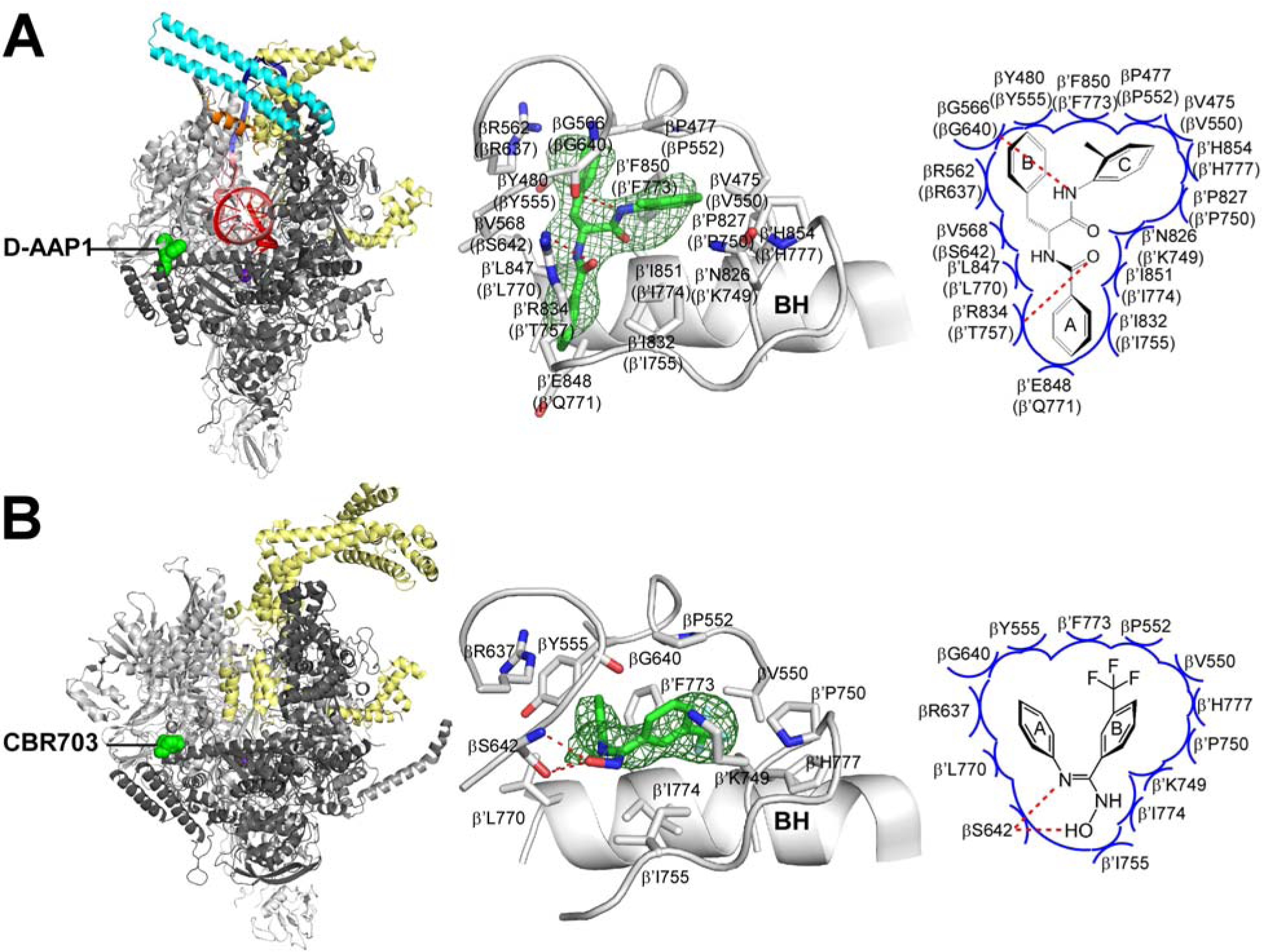
Relationship between binding sites of Mycobacterial-specific inhibitor D-AAP1 and Gram-negative-specific inhibitor CBR703: structures. (**A**) Structure of *Mtb* RPo in complex with D-AAP1. Left subpanel, overall structure. Middle subpanel, RNAP-inhibitor interactions. Right subpanel, schematic summary of RNAP-inhibitor interactions. Colors and labels as in Fig.3B. (**B**) Structure of *Eco* RNAP holoenzyme in complex with CBR703 (*17*). Subpanels, colors and labels as in (B).

**Fig. S12.**
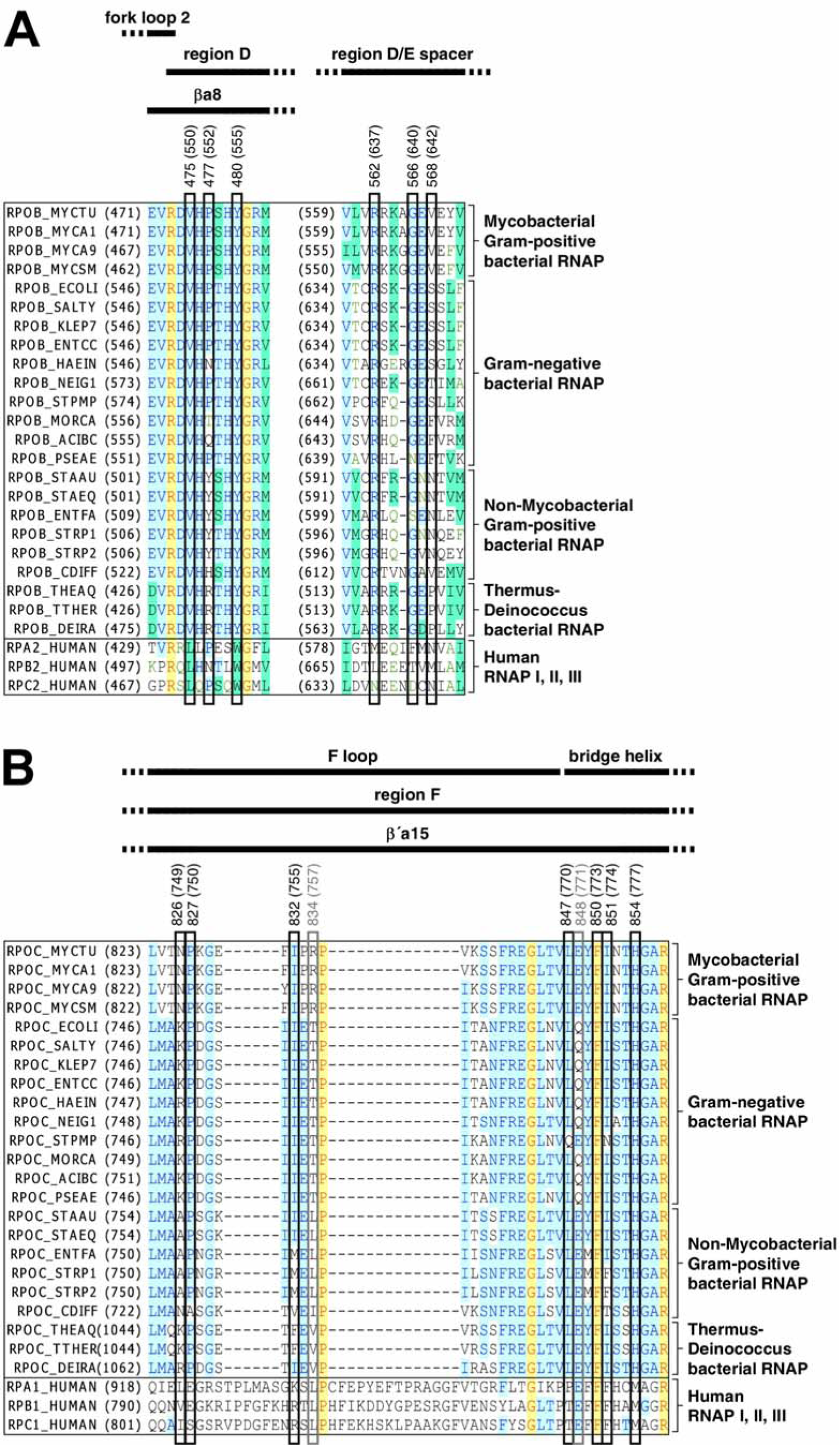
Relationship between binding sites of Mycobacterial-specific inhibitor D-AAP1 and Gram-negative-specific inhibitor CBR703: sequence alignments. Locations of residues that contact D-AAP1 and CBR703 in the sequences of RNAP β subunit (A) and RNAP β′ subunit (B). Sequence alignments for the β and β′ subunits of bacterial RNAP (top 23 sequences in each panel) and corresponding subunits of human RNAP I, RNAP II, and RNAP III (bottom three sequences in each panel), showing locations of RNAP residues that contact inhibitor in both *Mtb* RPo-D-AAP1 and *Eco* RNAP-CBR703 (black rectangles; identities from Fig. S11), RNAP residue that contacts inhibitor in *Mtb* RPo-D-AAP1 but not *Eco* RNAP-CBR703 (gray rectangle; identity from Fig. S11), locations of RNAP structural elements (top row of black bars; boundaries from *40-41*), and RNAP conserved regions (next two rows of black bars; boundaries from *8* and *42-43*). Species are as follows: *Mycobacterium tuberculosis* (MYCTU), *Mycobacterium avium* (MYCA1), *Mycobacterium abscessus* (MYCA9), *Mycobacterium smegmatis* (MYCSM), *Escherichia coli* (ECOLI), *Salmonella typhimurium* (SALTY), *Klebsiella pneumoniae* (KLEP7), *Enterococcus cloacae* (ENTCC), *Vibrio cholerae* (VIBCH), *Haemophilus influenzae* (HAEIN), Neisseria gonorrhoeae (NEIG1), Stenotrophomonas maltophilia (STPMP), Moraxella catarrhalis (MORCA), *Acinetobacter baumannii* (ACIBC), *Pseudomonas aeruginosa* (PSEAE), *Staphylococcus aureus* (STAAU), *Staphylococcus epidermidis* (STAEQ), *Enterococcus faecalis* (ENTFA), *Streptococcus pyogenes* (STRP1), *Streptococcus pneumoniae* (STRP2), *Clostridium difficile* (CDIFF), *Thermus thermophilus* (THETH), *Thermus aquaticus* (THEAQ), *Deinococcus radiodurans* (DEIRA), and *Homo sapiens* (HUMAN).

**Table S1.**
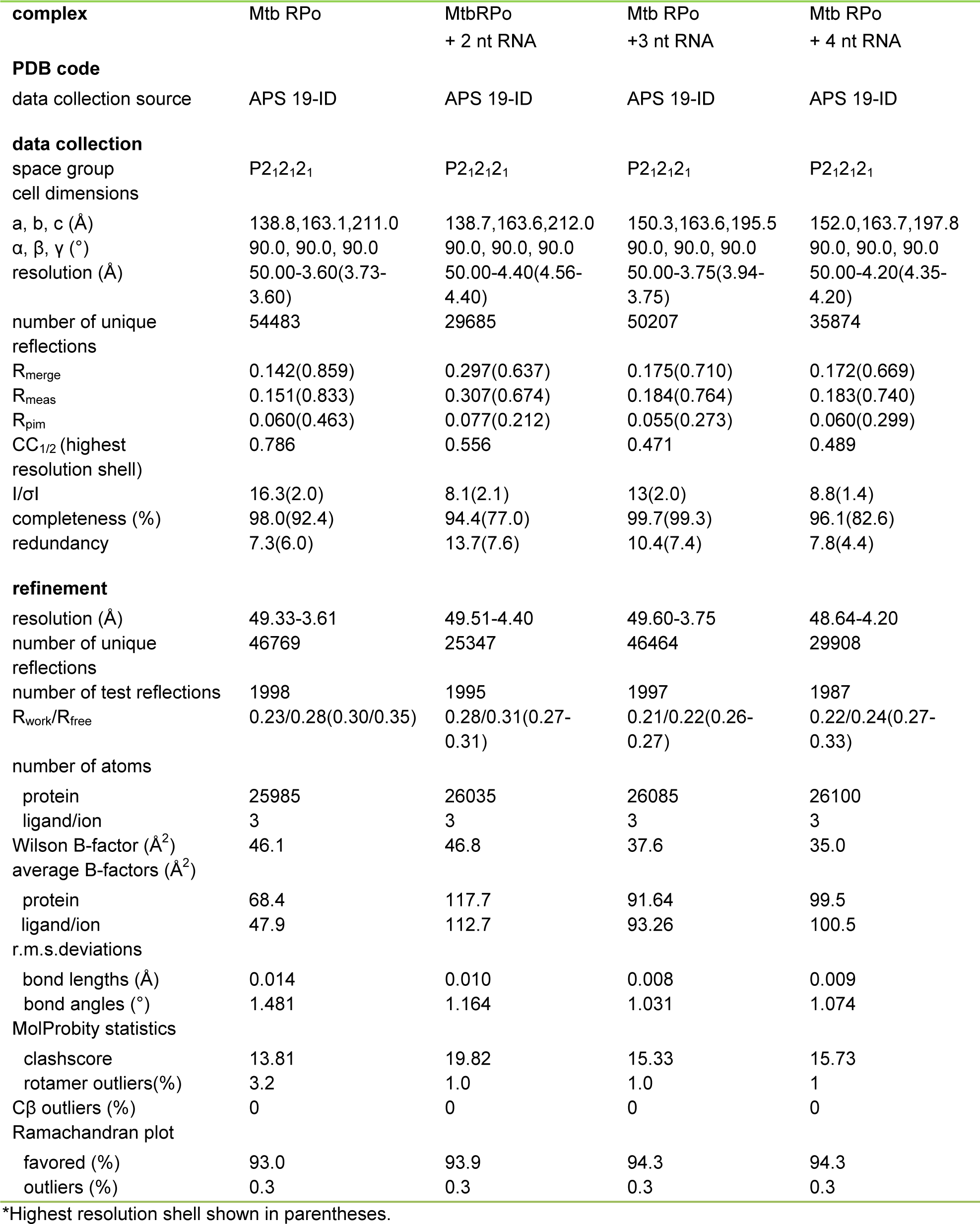
Structure data collection and refinement statistics.

| **complex** | Mtb RPo | MtbRPo<br>+ 2 nt RNA | Mtb RPo<br>+3 nt RNA | Mtb RPo<br>+ 4 nt RNA |
| --- | --- | --- | --- | --- |
| **PDB code** |  |  |  |  |
| data collection source | APS 19-ID | APS 19-ID | APS 19-ID | APS 19-ID |
| **data collection** |  |  |  |  |
| space group | P2<sub>1</sub>2<sub>1</sub>2<sub>1</sub> | P2<sub>1</sub>2<sub>1</sub>2<sub>1</sub> | P2<sub>1</sub>2<sub>1</sub>2<sub>1</sub> | P2<sub>1</sub>2<sub>1</sub>2<sub>1</sub> |
| cell dimensions |  |  |  |  |
| a, b, c (Å) | 138.8,163.1,211.0 | 138.7,163.6,212.0 | 150.3,163.6,195.5 | 152.0,163.7,197.8 |
| α, β, γ (°) | 90.0, 90.0, 90.0 | 90.0, 90.0, 90.0 | 90.0, 90.0, 90.0 | 90.0, 90.0, 90.0 |
| resolution (Å) | 50.00-3.60(3.73-3.60) | 50.00-4.40(4.56-4.40) | 50.00-3.75(3.94-3.75) | 50.00-4.20(4.35-4.20) |
| number of unique reflections | 54483 | 29685 | 50207 | 35874 |
| R<sub>merge</sub> | 0.142(0.859) | 0.297(0.637) | 0.175(0.710) | 0.172(0.669) |
| R<sub>meas</sub> | 0.151(0.833) | 0.307(0.674) | 0.184(0.764) | 0.183(0.740) |
| R<sub>pim</sub> | 0.060(0.463) | 0.077(0.212) | 0.055(0.273) | 0.060(0.299) |
| CC<sub>1/2</sub> (highest resolution shell) | 0.786 | 0.556 | 0.471 | 0.489 |
| I/σI | 16.3(2.0) | 8.1(2.1) | 13(2.0) | 8.8(1.4) |
| completeness (%) | 98.0(92.4) | 94.4(77.0) | 99.7(99.3) | 96.1(82.6) |
| redundancy | 7.3(6.0) | 13.7(7.6) | 10.4(7.4) | 7.8(4.4) |
| **refinement** |  |  |  |  |
| resolution (Å) | 49.33-3.61 | 49.51-4.40 | 49.60-3.75 | 48.64-4.20 |
| number of unique reflections | 46769 | 25347 | 46464 | 29908 |
| number of test reflections | 1998 | 1995 | 1997 | 1987 |
| R<sub>work</sub>/R<sub>free</sub> | 0.23/0.28(0.30/0.35) | 0.28/0.31(0.27-0.31) | 0.21/0.22(0.26-0.27) | 0.22/0.24(0.27-0.33) |
| number of atoms |  |  |  |  |
| protein | 25985 | 26035 | 26085 | 26100 |
| ligand/ion | 3 | 3 | 3 | 3 |
| Wilson B-factor (Å<sup>2</sup>) | 46.1 | 46.8 | 37.6 | 35.0 |
| average B-factors (Å<sup>2</sup>) |  |  |  |  |
| protein | 68.4 | 117.7 | 91.64 | 99.5 |
| ligand/ion | 47.9 | 112.7 | 93.26 | 100.5 |
| r.m.s.deviation |  |  |  |  |
| bond lengths (Å) | 0.014 | 0.010 | 0.008 | 0.009 |
| bond angles (°) | 1.481 | 1.164 | 1.031 | 1.074 |
| MolProbity statistics |  |  |  |  |
| clashscore | 13.81 | 19.82 | 15.33 | 15.73 |
| rotamer outliers(%) | 3.2 | 1.0 | 1.0 | 1 |
| Cβ outliers (%) | 0 | 0 | 0 | 0 |
| Ramachandran plot |  |  |  |  |
| favoured (%) | 93.0 | 93.9 | 94.3 | 94.3 |
| outliers (%) | 0.3 | 0.3 | 0.3 | 0.3 |

\*Highest resolution shell shown in parentheses.

**Table S2.** Structure data collection and refinement statistics.

| **complex** | *Mtb* RPo<br>+ Rif | *Mtb* RPo<br>+ Rif + 2 nt RNA | *Mtb* RPo<br>+ Rif + 3 nt RNA | *Mtb* RPo<br>+ Rif + 4nt RNA |
| --- | --- | --- | --- | --- |
| **PDB code** |  |  |  |  |
| data collection source | APS 19-ID | APS 19-ID | APS 19-ID | APS 19-ID |
| **data collection** |  |  |  |  |
| space group | P2<sub>1</sub>2<sub>1</sub>2<sub>1</sub> | P2<sub>1</sub>2<sub>1</sub>2<sub>1</sub> | P2<sub>1</sub>2<sub>1</sub>2<sub>1</sub> | P2<sub>1</sub>2<sub>1</sub>2<sub>1</sub> |
| cell dimensions |  |  |  |  |
| a, b, c (Å) | 154.4,164.2,200.2 | 153.2,164.9,199.9 | 150.1,167.4,195.2 | 152.1,163.1,197.9 |
| α, β, γ (°) | 90.0, 90.0, 90.0 | 90.0, 90.0, 90.0 | 90.0, 90.0, 90.0 | 90.0, 90.0, 90.0 |
| resolution (Å) | 50.00-4.25(4.40-4.25) | 50.00-3.80(3.94-3.80) | 50.00-3.80(3.94-3.80) | 50.00-4.00(4.14-4.00) |
| number of unique<br>reflections | 33,869 | 47,440 | 48,275 | 41,864 |
| R<sub>merge</sub> | 0.066(0.543) | 0.134(0.800) | 0.169(0.903) | 0.096(0.875) |
| R<sub>meas</sub> | 0.072(0.609) | 0.142(0.901) | 0.178(0.965) | 0.101(0.931) |
| R<sub>pim</sub> | 0.028(0.270) | 0.046(0.401) | 0.053(0.331) | 0.030(0.312) |
| CC<sub>1/2</sub> (highest<br>resolution shell) | 0.697 | 0.432 | 0.343 | 0.413 |
| I/σI | 20.2(2.0) | 14.2(1.6) | 11.0(2.0) | 23.5(1.9) |
| completeness (%) | 97.0(89.0) | 97.7(92.6) | 97.4(94.3) | 99.8(98.6) |
| redundancy | 5.9(4.5) | 7.5(4.3) | 9.9(7.5) | 11.1(8.2) |
| **refinement** |  |  |  |  |
| resolution (Å) | 49.78-4.29 | 50.09-3.84 | 48.81-3.80 | 49.48-4.00 |
| number of unique<br>reflections | 29562 | 39783 | 43854 | 38089 |
| number of test<br>reflections | 2005 | 1996 | 2000 | 1990 |
| R<sub>work</sub>/R<sub>free</sub> | 0.21/0.26(0.23-0.27) | 0.20/0.26(0.24/0.30) | 0.20/0.26(0.27-0.32) | 0.20/0.25(0.27-0.30) |
| number of atoms |  |  |  |  |
| protein | 26018 | 26035 | 26058 | 26080 |
| ligand/ion | 62 | 62 | 62 | 62 |
| Wilson B-factor (Å<sup>2</sup>) | 31.78 | 27.46 | 19.98 | 30.93 |
| average B-factors<br>(Å<sup>2</sup>) |  |  |  |  |
| protein | 86.9 | 89.4 | 90.8 | 88.4 |
| ligand/ion | 58.4 | 57.2 | 91.4 | 60.2 |
| r.m.s.deviation |  |  |  |  |
| bond lengths (Å) | 0.011 | 0.012 | 0.011 | 0.013 |
| bond angles (°) | 1.376 | 1.422 | 1.386 | 1.464 |
| MolProbity statistics |  |  |  |  |
| clashscore | 15.04 | 15.45 | 12.67 | 16.37 |
| rotamer outliers( %) | 2.8 | 3.2 | 2.9 | 3.3 |
| Cβ outliers (%) | 0 | 0 | 0 | 0 |
| Ramachandran plot |  |  |  |  |
| favored (%) | 92.3 | 92.0 | 93.1 | 92.3 |
| outliers (%) | 0.6 | 0.3 | 0.6 | 0.4 |

\*Highest resolution shell shown in parentheses.

**Table S3.** Structure data collection and refinement statistics.

| **complex** | *Mtb* RPo<br>+ AAP1 | *Mtb* RPo<br>+ D-IX336 | *Mtb* RPo<br>+ Rif + D-AAP1 | *Mtb* Se-β' *Mtb*SI |
| --- | --- | --- | --- | --- |
| **PDB code** |  |  |  |  |
| data collection source | APS 19-ID | APS 19-ID | APS 19-ID | APS 19-ID |
| **data collection** |  |  |  |  |
| space group | P2<sub>1</sub>2<sub>1</sub>2<sub>1</sub> | P2<sub>1</sub>2<sub>1</sub>2<sub>1</sub> | P2<sub>1</sub>2<sub>1</sub>2<sub>1</sub> | P2<sub>1</sub>2<sub>1</sub>2<sub>1</sub> |
| cell dimensions |  |  |  |  |
| a, b, c (Å) | 151.9, 164.6, 196.5 | 153.0, 164.6, 198.0 | 151.4, 162.1, 194.3 | 40.8, 97.5, 53.7 |
| $\alpha$, $\beta$, $\gamma$ (°) | 90.0, 90.0, 90.0 | 90.0, 90.0, 90.0 | 90.0, 90.0, 90.0 | 90.0, 93.3, 90.0 |
| resolution (Å) | 50.00-4.00(4.14-4.00) | 50.00-4.50(4.66-4.50) | 50.00-4.00(4.14-4.00) | 50.00-2.20(2.28-2.20) |
| number of unique reflections | 42228 | 30213 | 41653 | 21229 |
| R<sub>merge</sub> | 0.124(0.641) | 0.116(0.468) | 0.243(0.801) | 0.075(0.508) |
| R<sub>meas</sub> | 0.132(0.694) | 0.122(0.537) | 0.258(0.862) | 0.086(0.593) |
| R<sub>pim</sub> | 0.044(0.259) | 0.038(0.253) | 0.087(0.313) | 0.040(0.302) |
| CC<sub>1/2</sub> (highest resolution shell) | 0.554 | 0.525 | 0.490 | 0.791 |
| I/σI | 15.8(3.0) | 15.5(1.8) | 8.1(2.1) | 16.5(5.4) |
| completeness (%) | 99.7(98.4) | 98.2(93.2) | 99.8(99.9) | 99.4(96.8) |
| redundancy | 8.2(6.6) | 7.7(4.0) | 8.3(7.4) | 4.3(3.5) |
| **refinement** |  |  |  |  |
| resolution (Å) | 47.89-4.00 | 48.72-4.50 | 49.24-4.00 | 36.09-2.20 |
| number of unique reflections | 40742 | 26771 | 39399 | 18943 |
| number of test reflections | 1982 | 2712 | 2014 | 1835 |
| R<sub>work</sub>/R<sub>free</sub> | 0.21/0.27(0.23/0.27) | 0.21/0.23(0.25-0.27) | 0.21/0.26(0.28-0.34) | 0.21/0.26(0.30-0.34) |
| number of atoms |  |  |  |  |
| protein | 26080 | 26080 | 25966 | 2153 |
| ligand/ion | 30 | 29 | 89 |  |
| water |  |  |  | 45 |
| Wilson B-factor (Å<sup>2</sup>) | 31.47 | 37.18 | 40.67 | 44.21 |
| average B-factors (Å<sup>2</sup>) |  |  |  |  |
| protein | 33.9 | 93.0 | 122.3 | 62.6 |
| ligand/ion | 25.9 | 89.1 | 94.6 |  |
| r.m.s. deviations |  |  |  |  |
| bond lengths (Å) | 0.013 | 0.012 | 0.011 | 0.009 |
| bond angles (°) | 1.415 | 1.092 | 1.402 | 1.142 |
| MolProbity statistics |  |  |  |  |
| clashscore | 15.85 | 14.69 | 15.61 | 5.17 |
| rotamer outliers (%) | 3.2 | 1.0 | 2.7 | 0.0 |
| Cβ outliers (%) | 0 | 0 | 0 | 0 |
| Ramachandran plot |  |  |  |  |
| favored (%) | 92.0 | 94.5 | 91.8 | 97.7 |
| outliers (%) | 0.4 | 0.3 | 0.5 | 0.0 |

\*Highest resolution shell shown in parentheses.

## References

1. Global Tuberculosis Report (World Health Organization, Geneva, 2016).

2. D. Rothstein, Rifamycins, alone and in combination, Cold Spring Harb. Perspect. Med. 6 (2016).

3. P. Aristoff, G. Garcia, P. Kirchoff, H. Showalter, Rifamycins--obstacles and opportunities, Tuberculosis 90, 94 (2010).

4. J. Lee, S. Borukhov, Bacterial RNA polymerase-DNA interaction--the driving force of gene expression and the target for drug action, Front. Mol. Biosci. 3, 73 (2016).

5. K. Murakami, Structural biology of bacterial RNA polymerase, Biomolecules 5, 848 (2015).

6. E. Hubin, A. Fay, C. Xu, J. Bean, R. Saecker, M. Glickman, S. Darst, E. Campbell, Structure and function of the mycobacterial transcription initiation complex with the essential regulator RbpA, eLife, in press.

7. Y. Zhang, Y. Feng, S. Chatterjee, S. Tuske, M. Ho, E. Arnold, R.H. Ebright, Structural basis of transcription initiation, Science 338, 1076 (2012).

8. W. Lane, S. Darst, Molecular evolution of multisubunit RNA polymerases: sequence analysis., J. Mol. Biol. 395, 671 (2010).

9. V. Mekler, E. Kortkhonjia, J. Mukhopadhyay, J. Knight, A. Revyakin, A. Kapanidis, W. Niu, Y. Ebright, R. Levy, R.H. Ebright, Structural organization of bacterial RNA polymerase holoenzyme and the RNA polymerase-promoter open complex, Cell 108, 599 (2002).

10. B. Bae, E. Davis, D. Brown, E. Campbell, S. Wigneshweraraj, S. A. Darst, Phage T7 Gp2 inhibition of *Escherichia coli* RNA polymerase involves misappropriation of σ^70^ domain 1.1, Proc. Natl. Acad. Sci. USA 110, 19772 (2013).

11. E. Campbell, N. Korzheva, A. Mustaev, K. Murakami, S. Nair, A. Goldfarb, S. Darst, Structural mechanism for rifampicin inhibition of bacterial RNA polymerase, Cell 104, 901 (2001).

12. I. Artsimovitch, M. Vassylyeva, D. Svetlov, V. Svetlov, A. Perederina, N. Igarashi, N. Matsugaki, S. Wakatsuki, T. Tahirov, D. Vassylyev, Allosteric modulation of the RNA polymerase catalytic reaction is an essential component of transcription control by rifamycins, Cell 122, 351 (2005).

13. V. Molodtsov, I. Nawarathne, N. Scharf, P. Kirchhoff, H. Showalter, G. Garcia, K. Murakami, X-ray crystal structures of the *Escherichia coli* RNA polymerase in complex with benzoxazinorifamycins, J. Med. Chem. 56, 4758–4763 (2013).

14. A. Mustaev, E. Zaychikov, K. Severinov, M. Kashlev, A. Polyakov, V. Nikiforov, A. Goldfarb, Topology of the RNA polymerase active center probed by chimeric rifampicin-nucleotide compounds., Proc Natl Acad Sci USA. 91, 12036 (1994).

15. A. Feklistov, V. Mekler, Q. Jiang, L. Westblade, H. Irschik, R. Jansen, A. Mustaev, S. Darst, R.H. Ebright, Rifamycins do not function by allosteric modulation of binding of Mg^2^+ to the RNA polymerase active center, Proc. Natl. Acad. Sci. USA 105, 14820 (2008).

16. I. Artsimovitch, C. Chu, A. Lynch, R. Landick, A new class of bacterial RNA polymerase inhibitor affects nucleotide addition, Science 302, 650 (2003).

17. Y. Feng, D. Degen, X. Wang, M. Gigliotti, S. Liu, Y. Zhang, D. Das, T. Michalchuk, Y. Ebright, M. Talaue, N. Connell, R.H. Ebright, Structural basis of transcription inhibition by CBR hydroxamidines and CBR pyrazoles, Structure 23, 1470 (2015).

18. B. Bae, D. Nayak, A. Ray, A. Mustaev, R. Landick, S. Darst, CBR antimicrobials inhibit RNA polymerase via at least two bridge-helix cap-mediated effects on nucleotide addition, Proc. Natl. Acad. Sci. USA 112, E4178 (2015).

## Supplementary References

19. J. Jacques, S. Rodrigue, R. Brzezinski, L. Gaudreau, A recombinant Mycobacterium tuberculosis in vitro transcription system, FEMSMicrobiol Lett. 255, 140–147 (2006).

20. L. Stols, C. Millard, I. Dementieva, M. Donnelly, Production of selenomethionine-labeled proteins in two-liter plastic bottles for structure determination, J. Struct. Funct. Genomics. 5,: 95–102 (2004).

21. R. Banerjee, P. Rudra, R. Prajapati, S. Sengupta, J. Mukhopadhyay, Optimization of recombinant *Mycobacterium tuberculosis* RNA polymerase expression and purification, Tuberculosis 94, 397404 (2014).

22. T. Heyduk, Y. Ma, H. Tang, R.H. Ebright, Fluorescence anisotropy: rapid, quantitative assay for protein-DNA and protein-protein interaction, Meths. Enzymol. 274, 492 (1996).

23. Y. Zhang, D. Degen, M. Ho, E. Sineva, K. Ebright, Y. Ebright, V. Mekler, H. Vahedian-Movahed, Y. Feng, R. Yin, S. Tuske, H. Irschik, R. Jansen, S. Maffioli, S. Donadio, E. Arnold, R.H. Ebright, GE23077 binds to the RNA polymerase ‘i’ and ‘i+1’ sites and prevents the binding of initiating nucleotides, eLife 3, e02450 (2014).

24. A. Srivastava, M. Talaue, S. Liu, D. Degen, R.Y. Ebright, E. Sineva, A. Chakraborty, S. Druzhinin, S. Chatterjee, J. Mukhopadhyay, Y. Ebright, A. Zozula, J. Shen, S. Sengupta, R. Niedfeldt, C. Xin, T. Kaneko, H. Irschik, R. Jansen, S. Donadio, N. Connell, R.H. Ebright, New target for inhibition of bacterial RNA polymerase: “switch region,” Curr. Opin. Microbiol. 14, 532–543 (2011).

25. D. Degen, Y. Feng, Y. Zhang, K. Ebright, Y. Ebright, M. Gigliotti, H. Vahedian-Movahed, S. Mandal, M. Talaue, N. Connell, E. Arnold, W. Fenical, R.H. Ebright, Transcription inhibition by the depsipeptide antibiotic salinamide A, eLife 3, e02451 (2014).

26. L. Collins, S. Franzblau, Microplate Alamar Blue assay versus BACTEC 460 system for high-throughput screening of compounds against *Mycobacterium tuberculosis, Antimicrob*. Agents Chemother. 41, 1004–1009 (1997)

27. Clinical and Laboratory Standards Institute (CLSI/NCCLS), Methods for Dilution Antimicrobial Susceptibility Tests for Bacteria that Grow Aerobically; Approved Standard, Eighth Edition. CLIS Document M07-A8 (CLSI, Wayne, PA).(2009).

28. K. Murphy, K. Papavinasasundaram, C. Sassetti, Mycobacterial recombineering, Meths. Mol. Biol. 1285, 177–199 (2015).

29. A. Horrevorts, M. Michel, K. Kerrebijn, Antibiotic interaction: interpretation of fractional inhibitory and fractional bactericidal concentration indices, Eur. J. Clin. Microbiol. 6, 502–503 (1987).

30. R. White, R. M. Manduru, J. Bosso, Comparison of three different in vitro methods of detecting synergy: time-kill, checkerboard, and E test, Antimicrob. Agents Chemother. 40, 1914–1918 (1996).

31. J. Meletiadis, S. Pournaras, E. Roilides, T. Walsh, Defining fractional inhibitory concentration index cutoffs for additive interactions based on self-drug additive combinations, Monte Carlo simulation analysis, and in vitro-in vivo correlation data for antifungal drug combinations against Aspergillus fumigatus, Antimicrob. Agents Chemother. 54, 602–609 (2010).

32. W. Ma, G. Sandri, S. Sarkar, Analysis of the Luria-Delbrück distribution using discrete convolution powers, J. Appl. Probab. 29, 255–267 (1992).

33. S. Sarkar, W. Ma, G. Sandri, On fluctuation analysis: a new, simple and efficient method for computing the expected number of mutants, Genetica 85, 173–179 (1992).

34. B. Hall, C. Ma, P. Liang, and K. Singh, Fluctuation Analysis CalculatOR: a web tool for the determination of mutation rate using Luria-Delbrück fluctuation analysis, Bioinformatics 25, 1564–1565 (2009).

35. F. Stewart, D. Gordon, B. Levin, Fluctuation analysis: The probability distribution of the number of mutants under different conditions, Genetics. 124, 175–185 (1990).

36. X. Otwinowski, W. Minor, Processing of X-ray diffraction data collected in oscillation mode, Meths. Enzymol. 276, 307 (1997).

37. Collaborative Computational Project, The CCP4 suite: programs for protein crystallography, Acta Cryst. D 50, 760 (1994).

38. P. Emsley, B. Lohkamp, W. G. Scott, K. Cowtan, Features and development of Coot, Acta Cryst. D 66, 486 (2010).

39. P. Adams et al., PHENIX: a comprehensive Python-based system for macromolecular structure solution, Acta Cryst, D 66, 213 (2010).

40. R. Weinzierl, The nucleotide addition cycle of RNA polymerase is controlled by two molecular hinges in the bridge helix domain. BMC Biol. 8, 134 (2010).

41. P. Hein, R. Landick, The bridge helix coordinates movements of modules in RNA polymerase, BMC Biol. 8, 141 (2010).

42. D. Sweetser, M. Nonet, R. Young, Prokaryotic and eukaryotic RNA polymerases have homologous core subunits, Proc. Natl. Acad. Sci. USA 84, 1192–1196 (1987).

43. R. Jokerst, J. Weeks, W. Zehring, A. Greenleaf, Analysis of the gene encoding the largest subunit of RNA polymerase II in Drosophila, Mol. Gen. Genet. 215, 266–275 (1989).

